# SREBP regulates a cell autonomous triglyceride:glycogen metabolic switch in *Drosophila*

**DOI:** 10.1101/2024.05.13.593915

**Authors:** Rupali Ugrankar-Banerjee, Sonakshi Srivastava, Son Tran, Jade Bowerman, Blessy Paul, Lauren G. Zacharias, Thomas P. Mathews, Ralph J. DeBerardinis, W Mike Henne

## Abstract

Tissues store nutrients as triglyceride (TG) or glycogen at specific ratios, but how these reserves are sensed and balanced remains poorly understood. Here we show that suppression of the major *de novo* lipogenesis (DNL) enzyme fatty acid synthase FASN1 in the *Drosophila* fat body (FB) triggers a cell autonomous metabolic switch characterized by severe fat depletion and substantial glycogen accumulation. Despite their lipid deficiency, *Drosophila* develop normally into adults but exhibit shortened lifespans and impaired female fecundity. Mechanistically, we identify SREBP as a key mediator of the metabolic rewiring from TG to glycogen storage. We reveal this response is triggered by fatty acid deficiency resulting from DNL blockage but is attenuated by dietary fatty acid supplementation. Collectively, we propose a SREBP-dependent feedback loop where diminished fatty acid availability promotes a metabolic TG-to-glycogen energy shift in the *Drosophila* FB, supporting organismal development at the expense of reproductive success. These findings reveal an underappreciated role for SREBP in regulating the balance of fat and carbohydrate storage to maintain tissue and organismal homeostasis.

**Key findings:**

- Fat body-specific FASN1 loss leads to fat-depleted but viable *Drosophila* that complete their developmental lifecycle by rewiring energy metabolism to store glycogen instead of fat
- DNL inhibited *Drosophila* are viable, but display shortened lifespan and reduced female fecundity
- FASN1-deficient larvae functionally rely on glycogen synthesis and glycolysis
- Metabolic screening reveals a SREBP-dependent TG:glycogen metabolic switch in response to inhibition of *de novo* fatty acid biosynthesis, which is suppressed by addition of dietary fatty acids

## Introduction

Organisms must sense and allocate nutrients to metabolism, development, and reproduction to maintain homeostasis and populations^1,2^. Dietary carbohydrates are mobilized to glucose and catabolized through glycolysis, the tricarboxylic acid (TCA) cycle, and oxidative phosphorylation to yield carbon dioxide and ATP, the cellular energy currency^3–5^. Excess glucose can be converted into fatty acids and stored as triglycerides (TG) in lipid droplets (LD), or condensed to glycogen for energy utilization^3,4,6–8^. TG and glycogen reserves are balanced in a tissue and cell-dependent manner, and perturbations in their storage are linked to metabolic disorders including obesity, type 2 diabetes, cardiovascular disease, and cancers^4,5,9–11^. However, the metabolic decision-making that dictates how tissues invest in and balance fat and glycogen stores, and how these reserves are sensed and communicate to ensure cell and organismal homeostasis, remain pervasive questions deeply rooted in understanding and treating metabolic diseases^1^.

*Drosophila melanogaster* is a powerful model to interrogate metabolic imbalance and human metabolic diseases. *Drosophila* encode functional homologs to ∼75% of human disease genes, and key metabolic pathways such as insulin and TOR signaling are evolutionarily conserved in flies^10,12,13^. Like humans, *Drosophila* can become obese, lipodystrophic, or diabetic in response to caloric cues or genetic mutations^14–16^. Their primary nutrient storage tissue is the fat body (FB) which shares features of the mammalian adipose tissue and liver^17,18^. In feeding larvae, most dietary carbohydrates are converted to TG and stored in LDs within the FB. Glycogen is also present in the FB at lower concentrations, but its role is not fully clear. Glycogen is quickly mobilized during fasting, and recent studies indicate it may be important for maintaining hemolymph trehalose, the predominant circulating sugar in *Drosophila*, present at ∼100-times the level of glucose^12,16,19^.

TG stores in the FB are classically thought to support *Drosophila* development as both an energy and biomass resource during pupariation and metamorphosis^20–22^, which is characterized by large-scale developmental remodeling^23–25^. However, there is no consensus as to whether FB TG is necessary for this development. Paradoxically, lipodystrophic *Drosophila*, such as the *dLipin* mutant, can survive to the late pharate adult stage, and ∼40% become adult flies^26^. Similarly, no lethality is observed for *dSeipin* mutants that exhibit lipid storage and mobilization defects, and adults are viable and fertile^15,27,28^. Studies instead point to the blood-circulating sugar trehalose as an essential energy source for metamorphosis, raising questions as to how fat and carbohydrate reserves crosstalk and contribute to *Drosophila* development^21,29,30^.

TG in the FB is produced from two lipid pools: absorbed lipids from the blood by lipophorin trafficking, or via *de novo* lipogenesis (DNL) that requires fatty acid synthase (FASN)^9,31^. FASN is evolutionarily conserved and catalyzes the synthesis of long chain fatty acids from carbohydrate-derived malonyl-CoA, acetyl-CoA, and the electron donor nicotinamide adenine dinucleotide phosphate (NADPH)^32,33^. *Drosophila* encode three *FASN* genes with distinct tissue expressions. *FASN1/CG3523* is ubiquitously expressed with highest expression in the FB, whereas *FASN2/CG3524* and *FASN3/CG17374* are expressed primarily in the body wall, oenocytes, and hemocytes (*FASN2*) or oenocytes and reproductive system (*FASN3*)^24,34–36^. *Drosophila* hypomorphs for *FASN1* and *FASN2* exhibit reduced larval and adult TG, while null mutants of *FASN1* and *FASN3* display embryonic or early larval developmental arrest^24,34^. Fatty acid synthesis is thus crucial for the generation of energy reserves, cellular membranes, and protection from sugar toxicity^32,35^. However, how FASN-driven lipid synthesis is coordinated with carbohydrate metabolism to enable homeostasis and development remains incompletely understood.

Nutrient reserves must be sensed by cells to maintain energy homeostasis and adapt to nutrient shortages, and alterations in this are drivers of major metabolic diseases and cancers. Key nutrient sensing pathways include the TOR, AMPK, and insulin signaling systems, which monitor carbohydrate, lipid, and amino acids, as well as energy reserves to coordinate organismal development^36–40^. Notably, mammalian isoforms of the sterol regulatory element-binding protein (SREBP), an insulin- and TOR-responsive transcription factor best characterized for its role in lipogenesis, have also been implicated in the regulation of carbohydrate metabolism. For example in the liver, SREBP-1c has been reported as required for insulin-dependent expression of glucokinase, a key glycolysis enzyme that catalyzes glucose phosphorylation to glucose-6-phosphate, a precursor for glycogen synthesis^41–43^. Furthermore, glycogen synthase kinase 3 (GSK3), a negative regulator of glycogen synthase (GlyS), inhibits SREBP-1c transcriptional activity in hepatocytes and adipocytes^44^. These observations raise the possibility that *SREBP-dependent transcriptional programs may contribute to the coordination of both lipid and carbohydrate metabolism*, although the underlying mechanisms remain poorly understood.

To address this knowledge gap in TG:glycogen crosstalk, here we dissect molecular players that orchestrate this energy balance by genetically targeting the major fatty acid synthase homolog FASN1 in the *Drosophila* FB. We find that FASN1 depletion, or inhibition of other early steps of DNL in the FB, leads to drastic TG loss accompanied by a marked boost in glycogen storage, suggesting a shift towards glycogen metabolism as an alternative energy storage pathway. These fat-depleted *Drosophila* are viable but short-lived and females are infertile. Mechanistically, we identify a cell-autonomous, SREBP-dependent TG:glycogen feedback switch that mediates this metabolic rewiring. We find impaired fatty acid biosynthesis promotes SREBP activation and triggers this response, which can be attenuated by dietary lipid supplementation. Collectively, we propose that *Drosophila* SREBP coordinates lipid and carbohydrate metabolism in the FB to adapt to fatty acid deficiency, thereby enabling organismal development at the expense of reproductive success.

## Results

### Fat body specific *FASN1* loss leads to severe lipid depletion in *Drosophila*

The fat body (FB) is the central metabolic organ in *Drosophila,* and represents much of the larval biomass necessary to achieve metamorphosis^18,20–22^. The FB lipid mass is composed of lipid droplet (LD) triglycerides (TGs), 90% of which is produced from *de novo* lipogenesis (DNL)^9,31,45^. To genetically manipulate these fat stores and investigate how they influence *Drosophila* development and metabolism, we utilized the FB-specific driver *Dcg-Gal4*^16,46^ to selectively knockdown (KD) the major fatty acid synthase *FASN1* (**Figure 1A,B**), which we previously identified as the predominant FASN paralog contributing to *de novo lipogenesis* (DNL) for TG synthesis in the FB^31^ (**SFigure 1A**). We confirmed that the FB-targeted FASN1 RNAi depletion significantly reduced *FASN1* mRNA transcripts (**Figure 1B**). Notably, the FB tissue remained easily detectable in *FASN1^FB-RNAi^* larvae and could be isolated for analysis. FBs dissected from *FASN1^FB-RNAi^*(*Dcg-Gal4>FASN1-RNAi*) *Drosophila* third instar (L3) late-feeding larvae (∼100-108 h After Egg Laying) sank in PBS buffer, indicating they lost buoyancy normally conferred by lipids, which was retained in RNAi control FBs, expressing RNAi against *mCherry* (*Dcg-Gal4>mCherry-RNAi*, hereafter referred to as *Dcg-Gal4* controls) (**SFigure 1B**). Despite this marked lipid depletion, *FASN1^FB-RNAi^* adipocytes were not reduced in size but instead retained their typical morphology and were even slightly larger than *Dcg-Gal4* control adipocytes (**Figure 1C,D**).

**Figure 1:**
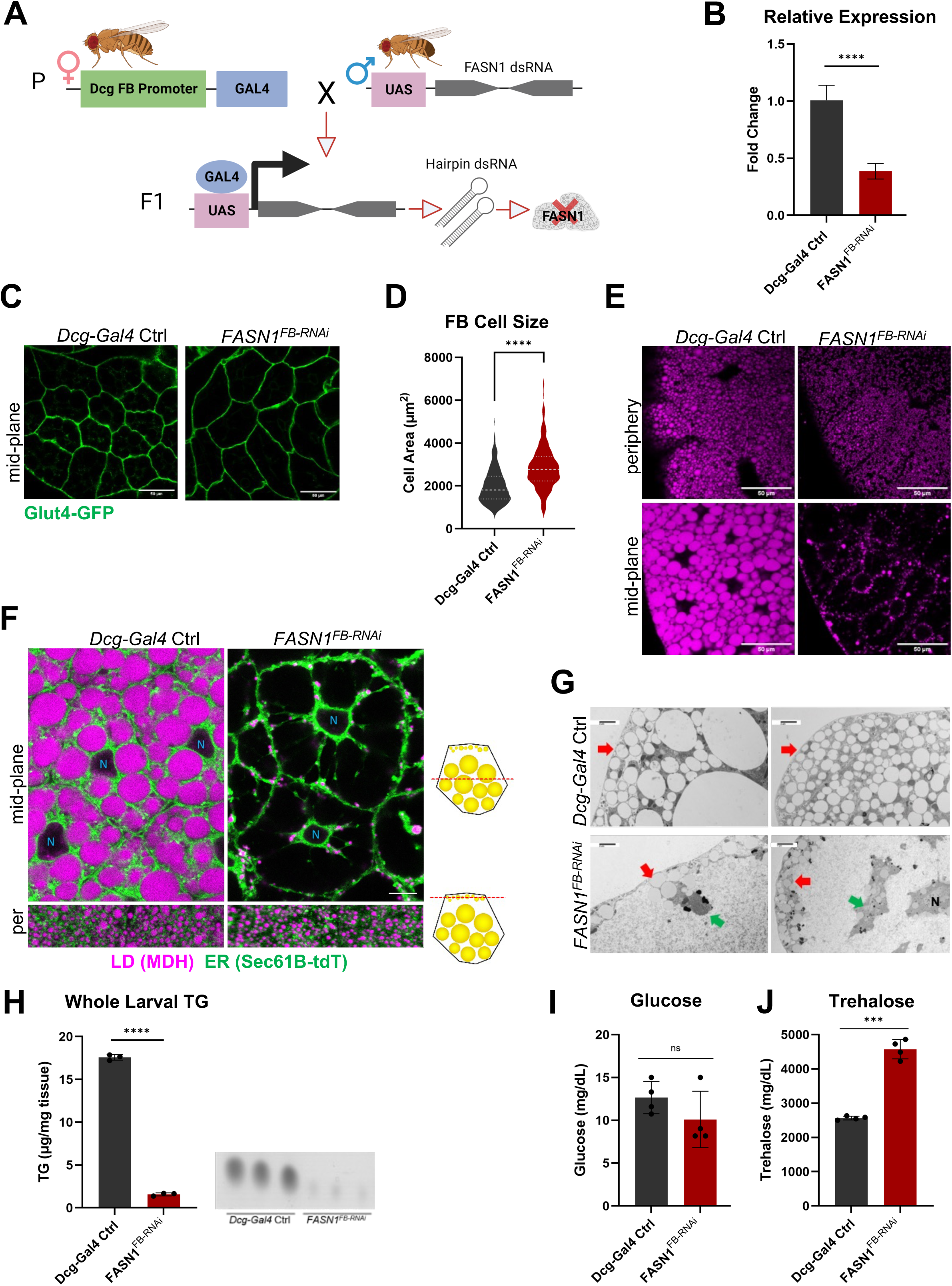
Loss of FASN1 in *Drosophila* fat body produces virtually fat less flies A) Cartoon of RNA interference-mediated KD using the UAS/Gal4 system; parental cross between females carrying tissue (fat body)-specific *Dcg-Gal4* driver and males containing the *UAS-FASN1 dsRNA* transgene generates progeny lacking *FASN1* in the FB. B) Relative expression of *FASN1* mRNA in *Dcg-Gal4>mCherry^RNAi^* (*Dcg-Gal4* control) and *Dcg-Gal4>UAS-FASN1^RNAi^* (*FASN1^FB-RNAi^*) FBs. Data are shown as mean ± SD from 3 biological replicates per genotype (FBs pooled from 20-30 late-feeding third instar L3 larvae per replicate). P values were determined using an unpaired, two-tailed Student’s t-test. C) Confocal images of *Dcg-Gal4* control and *FASN1^FB-RNAi^* FBs from L3 larvae expressing a membrane-associated reporter (*Dcg-Gal4>UAS-Glut4-GFP*) to visualize cell boundaries. Scale bar 50 µm. D) Violin plots of FB cell area measurements from 250 *Dcg-Gal4* control and 290 *FASN1^FB-RNAi^* adipocytes sampled in 19 and 32 fat body (FB) tissues, respectively. Statistical analyses were performed using mean cell area calculated for each tissue as the experimental unit. P values were determined using an unpaired, two-tailed Welch’s t-test. E) Confocal images of *Dcg-Gal4* control and *FASN1^FB-RNAi^* larval (L3) FB periphery and mid-plane sections stained with LD stain monodansylpentane (MDH). Scale bar 50 µm. F) Confocal images of *Dcg-Gal4* control and *FASN1^FB-RNAi^*FBs at their mid-plane and periphery from larvae expressing ER marker (*Dcg-Gal4>UAS-Sec61B-tDTomato*) and stained with MDH. *FASN1* loss results in adipocytes empty of LDs. Scale bar 10 µm. G) TEM micrographs of *Dcg-Gal4* control and *FASN1^FB-RNAi^* FB cells showing peripheral LDs (pLDs, red arrows). *FASN1^FB-RNAi^* cytoplasm appears largely vacant apart from small ‘islands’ of ER, LDs, and other organelles (green arrows). Scale bar 2-4 µm. H) Total triglycerides (TG) levels measured by thin layer chromatography (TLC) in whole *Dcg-Gal4* control and *FASN1^FB-RNAi^* L3 larvae. TLC plate representing TG bands corresponding to the quantified samples is shown alongside the graph (sample volumes loaded onto the plate were normalized to larval weight such that band intensities reflect TG abundance). Data are shown as mean ± SD from 3 biological replicates per genotype (10 larvae per replicate). P values were determined using an unpaired, two-tailed Welch’s t-test. I) *Dcg-Gal4* control and *FASN1^FB-RNAi^* larval hemolymph glucose levels (mg/DL). Data are shown as mean ± SD from 4 biological replicates per genotype (hemolymph pooled from 10-12 L3 larvae per replicate). P values were determined using an unpaired, two-tailed Welch’s t-test. J) *Dcg-Gal4* control and *FASN1^FB-RNAi^* larval hemolymph trehalose levels (mg/DL). Data are shown as mean ± SD from 4 biological replicates per genotype (hemolymph pooled from 10-12 L3 larvae per replicate). P values were determined using an unpaired, two-tailed Welch’s t-test.

To determine how FASN1 deficiency impacted LDs in the FB, we conducted z-section confocal imaging of FBs excised from larvae. *FASN1^FB-RNAi^*FBs displayed a near complete lack of large LDs normally visible in the tissue mid-plane (mid-plane LDs, or mLDs), which constitute the bulk of TG stores in FB cells^47^. However, *FASN1^FB-RNAi^* FBs did exhibit small peripheral LDs (pLDs) at the tissue surface, consistent with previous work indicating these pLDs are largely independent of DNL, and may potentially be supported by extracellular lipids delivered to the FB^31^ (**Figure 1E,F**). pLDs near the cell surface of larval FBs were also observed by transmission electron microscopy (TEM) (**Figure 1G, pLDs at red arrows**). In contrast, the cytoplasm of *FASN1^FB-RNAi^*FB cells was largely devoid of mLDs and appeared mostly vacant, containing small ‘islands’ of ER, a few residual LDs, and other organelles (**Figure 1G, green arrows**). As expected, total larval TG was significantly reduced by ∼85% in *FASN1* KD compared to control animals (**Figure 1H**). As expected, FB-specific KD of *FASN3*, and a global hypomorphic mutation of *FASN2*, did not produce a pronounced depletion of FB LDs, consistent with the limited roles of these *FASN* paralogs in FB lipid storage^47^ (**SFigure 1C, see Table S1 for list of *Drosophila* stocks**).

To further validate that TG loss in the FB was the result of FB-specific FASN1 depletion, we also targeted FASN1 using two additional FB drivers, *Cg-Gal4* and *r4-Gal4*, both of which also resulted in LD depletion akin to that observed with *Dcg-Gal4* (**SFigure 1D**). Perturbations in FB fat storage can sometimes cause ectopic lipid accumulation in other tissues such as the mid-gut^48^, however gut TG levels were 50% lower in *FASN1^FB-RNAi^* larvae relative to controls (**SFigure 1E**). These observations collectively suggested that FB-specific *FASN1* depletion led to the loss of the majority of larval TG stores.

Since lipogenesis is essential to protect the animal from glucotoxicity^16,18^ we also examined how the disruption of dietary carbohydrate-to-fat conversion in *FASN1^FB-RNAi^* larvae impacted blood (i.e. hemolymph) circulating glucose and trehalose. We measured these in control and *FASN1^FB-RNAi^*L3 late-feeding larvae. Notably, circulating glucose levels were not significantly altered, but total circulating carbohydrates comprised primarily of trehalose were substantially up in *FASN1^FB-RNAi^* larvae, suggesting they were hypertrehalosemic (**Figure 1I,J**). Collectively, this indicates that FB-targeted reduction of FASN1 results in a substantial depletion of larval fat reserves and elevated circulating carbohydrates.

### *FASN1^FB-RNAi^* flies are viable but exhibit shortened lifespan, starvation sensitivity, and reduced fecundity

Despite their severely impaired lipid storage and hypertrehalosemic ‘diabetic-like’ state^16^, *FASN1^FB-RNAi^* larvae appeared healthy, exhibited developmental progression similar to that of age-matched *Dcg-Gal4* controls, and attained full larval body size (**Figure 2A, top panel**). In L3 late-feeding larvae, the FB tissue visualized by GFP fluorescence from a *UAS-GFP* transgene expressed in the *Dcg-Gal4* control and *FASN1^FB-RNAi^* backgrounds (**Figure 2A, bottom panel, yellow arrows**), appeared normal in size. However, the *FASN1*-depleted FB appeared notably translucent, consistent with reduced lipid storage, in contrast to the opaque white characteristic of fat-engorged control tissue (**Figure 2A, yellow arrows**). Remarkably *FASN1^FB-RNAi^* larvae successfully pupated, giving rise to pupae that were similar in length to controls though very slightly narrower (**Figure 2B,C**). Moreover, they completed metamorphosis and eclosed at rates only slightly lower to those of control flies (87% vs. 79%, **Figure 2D**), emerging as normal-sized adults with body weights comparable to controls (**Figure 2E,F**). As expected, *FASN1^FB-RNAi^* 7-10 days old female and male flies have virtually undetectable TG (**Figure 2G**), and an adult fat body was not easily discernable in these flies. Given this dramatic fat deficiency, we investigated how this influenced *FASN1^FB-RNAi^*fly lifespan and metabolic stress responses. We found *FASN1^FB-RNAi^*flies of both sexes displayed significantly shortened median lifespan on standard food (**Figure 2H**). Moreover, *FASN1^FB-RNAi^*adults died roughly twice as quickly when maintained on water but no food source, indicating starvation sensitivity consistent with a lack of TG reserves (**Figure 2I**).

**Figure 2:**
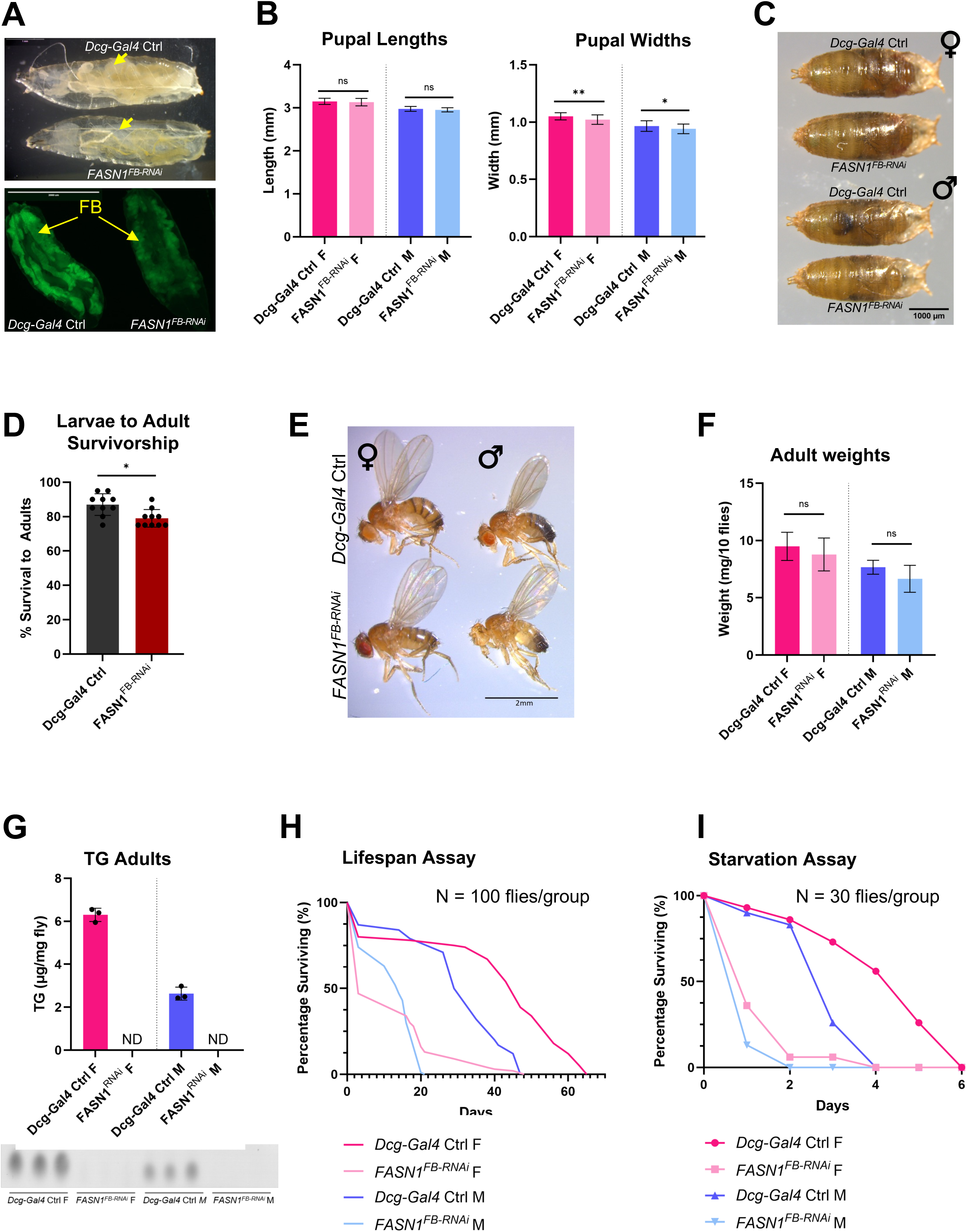
*FASN1*-deficient flies are viable but exhibit reduced lifespan and increased starvation sensitivity A) (Top panel) *Dcg-Gal4* control and *FASN1^FB-RNAi^*age-matched late-feeding third instar (L3) larvae. Scale bar 1 mm. (Bottom panel) Fluorescence micrograph of *Dcg-Gal4* control and *FASN1^FB-RNAi^* larvae expressing soluble GFP in their FBs (*Dcg-Gal4>UAS-GFP*). Scale bar 2 mm. FBs indicated by yellow arrows in both panels. B) Pupal sizes (Length = L, Width = W) for *Dcg-Gal4* control and *FASN1^FB-RNAi^* pupae, measured using digital Vernier calipers. Data are shown as mean ± SD of 30 male and 30 female pharate pupae per genotype, analyzed separately. P values were determined using an unpaired, two-tailed Welch’s t-test. C) Visual comparison of *Dcg-Gal4* control and *FASN1^FB-RNAi^* female (top) and male (bottom) pharate pupae. Scale bar 1000 µm. D) Percentage of *Dcg-Gal4* control and *FASN1^FB-RNAi^* L3 larvae successfully completing development to adult flies. Data are shown as mean ± SD from 10 independent experiments (200 L3 larvae per genotype across 10 experimental repeats). P values were determined using a two-tailed Fisher’s exact test on pooled eclosion counts. E) Visual comparison of *Dcg-Gal4* control and *FASN1^FB-RNAi^* female (left) and male (right) adult flies. Scale bar 2 mm. F) Weights of female and male *Dcg-Gal4* control and *FASN1^FB-RNAi^* adult flies, aged 7-10 days. Data are shown as mean ± SD from 9 biological replicates per genotype, with males and females weighed separately in groups of 10 flies. P values were determined using an unpaired, two-tailed Welch’s t-test. G) Total triglycerides (TG) measured by thin layer chromatography (TLC) in *Dcg-Gal4* control and *FASN1^FB-RNAi^* 7-10 days old female and male flies. TLC plate representing TG bands corresponding to the quantified samples is shown below the graph (sample volumes were normalized to fly weight). Data are shown as mean ± SD from 3 biological replicates per genotype (10 flies per replicate). P values were determined using an unpaired, two-tailed Welch’s t-test. H) Longevity assay performed for *Dcg-Gal4* control and *FASN1^FB-RNAi^* adult females and males (100 male and 100 female flies per genotype). I) Starvation assay using 7-10 days old *Dcg-Gal4* control and *FASN1^FB-RNAi^* female and male flies (30 males and 30 female flies per genotype).

As nutrient reserves are tightly linked with reproductive success^49,50^, we next examined how early DNL blockage in the FB impacted fecundity by monitoring whether *FASN1^FB-RNAi^* female and male flies could successfully generate progeny. Indeed, eggs deposited by control (*Dcg-Gal4)* females mated with *FASN1^FB-RNAi^* males showed slightly decreased viability relative to those from *Dcg-Gal4* fathers, suggesting *FASN1* loss in the male FB mildly perturbed fecundity (S**Figure 2A**). Strikingly, *FASN1^FB-RNAi^*females mated with either control or *FASN1^FB-RNAi^* males failed to lay eggs, indicating *FASN1^FB-RNAi^* females exhibited substantial infertility.

The *Drosophila* female fly reproductive system consists of two ovaries, each containing 20-30 ovarioles, which are organized as linear progressions of maturing egg chambers^51,52^. Dissection of the internal reproductive organs of *FASN1^FB-RNAi^* females revealed shrunken and underdeveloped ovaries that did not produce mature oocytes (**SFigure 2B,C, oocytes denoted by white arrows**). Close examination revealed *FASN1^FB-RNAi^* egg chambers were arrested at stage ∼9-10, when follicle cells normally migrate to surround the oocyte (**SFigure 2D, follicle cells denoted by yellow arrows, oocyte by white arrows**). In control ovaries, LDs accumulated in nurse cells (**SFigure 2E, denoted by orange arrows**) and were subsequently transferred to the growing oocyte (**SFigure 2E, denoted by purple arrow**). In contrast, *FASN1^FB-RNAi^* egg chambers were largely devoid of LDs, consistent with reports their deficiency correlates with defective oogenesis^51–54^. Failure to pass this key metabolic checkpoint in oocyte maturation likely resulted in degeneration of the egg chambers, as no discernable cells with nuclear DAPI staining were visible in the interior of *FASN1^FB-RNAi^* ovaries (**SFigure 2F, blue arrows).** Collectively, this indicates that *FASN1^FB-RNAi^ Drosophila* larvae, despite lacking a majority of their FB TG stores, are still able to achieve the critical weight required for pupation and metamorphosis. However, while FASN1-dependent DNL in the FB is dispensable for animal development, FASN1 loss compromises lifespan, starvation resistance, and female fecundity.

### FASN1-deficient *Drosophila* FBs metabolically rely on glycogen storage for development

The fact that perturbation of an early step in DNL (i.e. FASN1 depletion) in the FB led to viable but largely fat depleted *Drosophila* provided a unique opportunity to dissect how adipose metabolism may be rewired to support animal development in such conditions. *Drosophila* FB lipids are proposed to play crucial roles in development, serving as an energy source for metamorphosis, biomaterials for growth, and a metabolic buffer to counter caloric overload^10,22,24^. Given this, we investigated what proteins and pathways were necessary for metamorphosis in *FASN1^FB-RNAi^* animals. We conducted unbiased global liquid chromatography tandem mass spectrometry (LC-MS/MS) proteomics of isolated FBs from control and *FASN1^FB-RNAi^*L3 late-feeding larvae. Principal component analysis (PCA) revealed control and *FASN1^FB-RNAi^* FBs exhibited substantial differences in the relative abundances of many proteins (**Figure 3A, Table S2**). Proteomics also confirmed efficient *FASN1* KD, as FASN1 was among the most strongly downregulated proteins in *FASN1^FB-RNAi^*FBs (**Figure 3B**). We also noted reduced abundances of proteins important for TG synthesis or storage, such as glycerol kinase Gk1, perilipins Lsd-1 and Lsd-2, proteins involved in TG breakdown such as Brummer (Bmm, a mammalian ATGL homolog), as well as pdgy and CPT2 (both involved in fatty acid β-oxidation) (**Figure 3B**). In line with this, Gene Ontology (GO) term pathway analysis revealed lipid metabolic processes, particularly those related to fatty acid, TG biosynthesis, and fatty acid catabolism, were among the most decreased in *FASN1^FB-RNAi^* FBs (**Figure 3C**). Conversely, expression of upstream lipogenic genes, including ATP-citrate lyase (ATPCL) and Acetyl CoA carboxylase (ACC), were increased in FASN1-depleted FB (**Figure 3B**).

**Figure 3:**
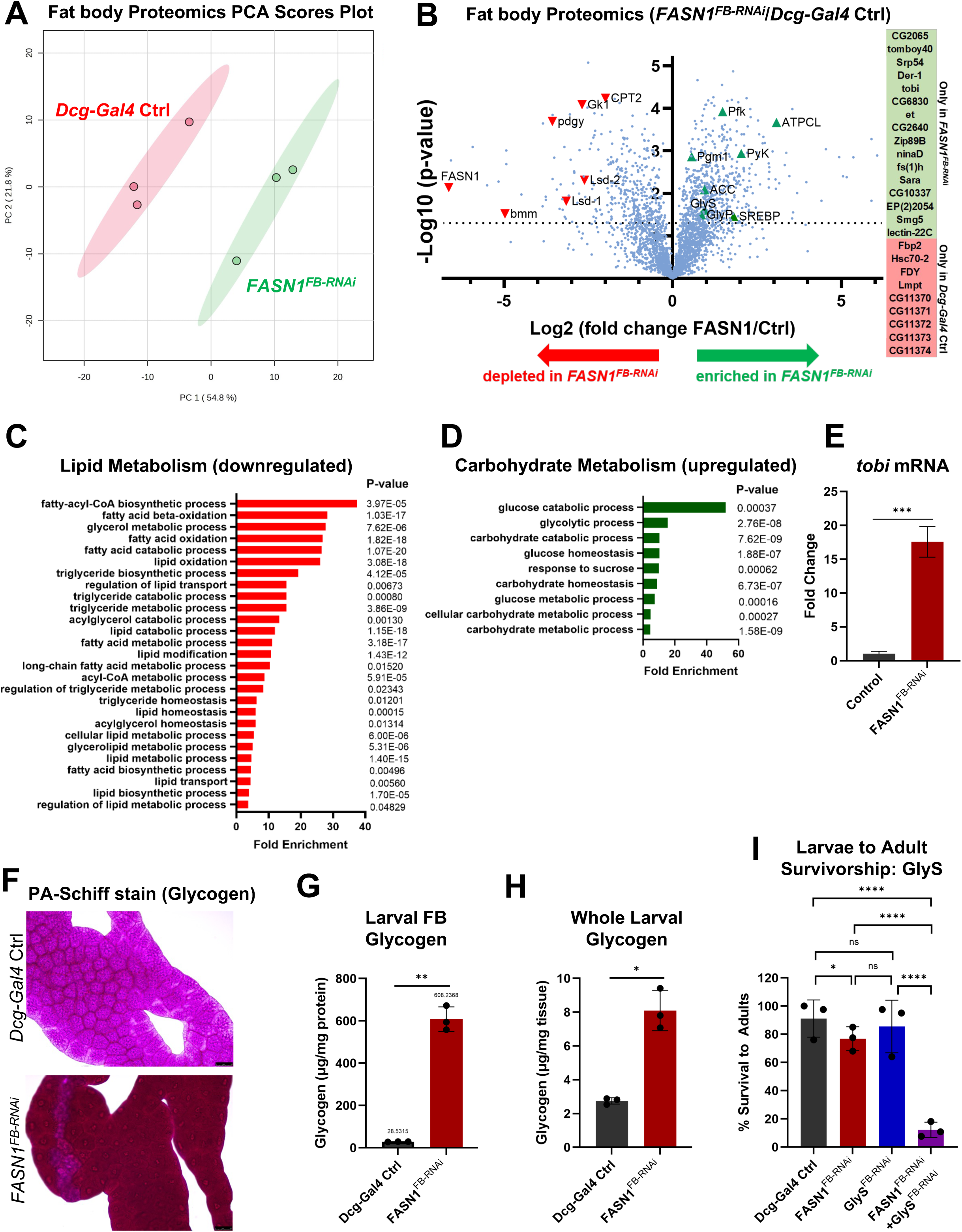
*FASN1*-deficient larvae channel dietary glucose to glycogen as energy reserves A) Principal Component Analysis (PCA) for global proteomics performed in FBs isolated from *Dcg-Gal4* control and *FASN1^FB-RNAi^* L3 larvae. B) Volcano plot of proteins detected by LC-MS/MS analyses performed in FBs isolated from *Dcg-Gal4* control and *FASN1^FB-RNAi^* L3 larvae. C) Lipid metabolism processes were revealed to be amongst the most downregulated in *FASN1^FB-RNAi^* FBs. GO enrichment analysis of proteomics data was performed using Pangea. D) Carbohydrates metabolism processes downregulated in *FASN1^FB-RNAi^* FBs (Pangea). E) Relative expression of *tobi* mRNA in isolated FBs from *Dcg-Gal4* control and *FASN1^FB-RNAi^*larvae, measured by qPCR. Data are shown as mean ± SD from 3 biological replicates per genotype (FBs pooled from 20-30 L3 larvae per replicate). P values were determined using an unpaired, two-tailed Student’s t-test. F) Periodic acid-Schiff (PAS) glycogen stain of larval FBs from *Dcg-Gal4* control and *FASN1^FB-RNAi^* L3 larvae. Scale bar 50 µm. G) Biochemical quantification of glycogen extracted from *Dcg-Gal4* control and *FASN1^FB-RNAi^* larval FBs. Data are shown as mean ± SD from 3 biological replicates per genotype (FBs from 25-20 L3 larvae per replicate), normalized to total protein. P values were determined using an unpaired, two-tailed Welch’s t-test. H) Biochemical quantification of glycogen extracted from *Dcg-Gal4* control and *FASN1^FB-RNAi^* whole L3 larvae. Data are shown as mean ± SD from 3 biological replicates per genotype (15 larvae per replicate), normalized to weight. P values were determined using an unpaired, two-tailed Welch’s t-test. I) Percentage of L3 larvae (*Dcg-Gal4* control, *FASN1^FB-RNAi^*, *GlyS^FB-RNAi^*, *GlyS^FB-RNAi^*+ *FASN1^FB-RNAi^*) successfully completing metamorphosis to yield adult flies. Data are shown as mean ± SD from 3 independent experiments (110 L3 larvae per genotype across 3 experimental repeats). P values were determined using a two-tailed Fisher’s exact test on pooled eclosion counts.

Whereas lipid metabolism generally appeared blunted in *FASN1^FB-RNAi^*FBs, GO term analysis indicated carbohydrate catabolic and glycolytic processes were among the most upregulated (**Figure 3D**). Notable among the upregulated glycolytic genes were phosphofructokinase (Pfk), which catalyzes the committed step of glycolysis by generating fructose-1,6-bisphosphate, and pyruvate kinase (Pyk), which catalyzes the final step of glycolysis to produce pyruvate, a precursor of acetyl-CoA. Among the most elevated carbohydrate-related proteins was target-of-brain-insulin (tobi), a carbohydrate metabolism associated protein^55,56^, which was only detected in FBs from *FASN1^FB-RNAi^* larvae (**Figure 3B, green box**). Consistent with this, *tobi* mRNA levels were elevated nearly 20-fold in *FASN1^FB-RNAi^* FBs relative to controls (**Figure 3E).** Tobi is a conserved putative alpha-glucosidase expressed primarily in FB and gut, and is activated during sugar mobilization^55,56^. Alpha-glucosidases like tobi are speculated to contribute to glycogen breakdown. Given these signatures of carbohydrate metabolism, we hypothesized carbohydrate storage in the form of glycogen may be altered in *FASN1^FB-RNAi^*FBs. Therefore, we examined FB glycogen levels in control and *FASN1^FB-RNAi^* FBs. Strikingly, periodic acid-Schiff (PAS) staining revealed a marked boost in glycogen accumulation in *FASN1^FB-RNAi^* FBs compared with controls (**Figure 3F**). Biochemical glycogen measurements confirmed a remarkable ∼20-fold increase in glycogen levels in the FB, accompanied by an ∼3-fold glycogen increase in whole animal glycogen; this difference likely reflects glycogen storage in other tissues, including muscle, a major glycogen reservoir^57^ (**Figure 3G,H).** Although glucose derived from gut contents may contribute to whole-animal glucose measurements, food intake, estimated by quantifying Coomassie dye in the guts of late feeding L3 larvae cultured on dye-supplemented food, was comparable between *Dcg-Gal4* controls and *FASN1^FB-RNAi^* larvae. This suggested that differences in food consumption are unlikely to contribute to the elevated whole-animal glycogen levels in FASN1-KD larvae (**SFigure 3A**).

In line with this boosted carbohydrate storage in *FASN1^FB-RNAi^* larvae, qPCR revealed induction of key enzymes involved in glycogen synthesis (glycogenesis) and glycogen breakdown (glycogenolysis) (**SFigure 3B**). In particular glycogen synthase (GlyS) and glycogen phosphorylase (GlyP) are attributed to glycogen synthesis and breakdown in the *Drosophila* FB^57^. This collectively suggested that in the absence of fat stores, larvae may channel nutrients into FB glycogen for energy storage and utilization. From this, we hypothesized that *FASN1^FB-RNAi^*larvae depended on carbohydrate metabolism for development.

To determine whether these glycogen stores were necessary for development, we RNAi depleted the glycogen synthesis enzyme GlyS in control and *FASN1^FB-RNAi^* animals. Intriguingly, whereas RNAi loss of GlyS alone did not impact larval development into flies, loss of GlyS in combination with *FASN1^FB-RNAi^* led to developmental arrest and lethality during the pupal stage, with only a few flies reaching adulthood (**Figure 3I**). This indicated that glycogen synthesis was essential for adult *Drosophila* survival when FASN1 was depleted. Next, we monitored glycogen levels in control and *FASN1^FB-RNAi^* animals during metamorphosis. In controls, glycogen levels remained steady throughout metamorphosis (**SFigure 3C**). In contrast, glycogen levels in *FASN1^FB-RNAi^* animals declined rapidly during the transition from late feeding larvae to pharate adults (abbreviated PA) **(SFigure 3C**). In fact, *FASN1^FB-RNAi^* pharate adults contained only ∼15% of their initial glycogen stores, consistent with FB glycogen mobilization during metamorphosis to support development. Conversely, controls exhibited a pronounced decline in TG by ∼40%, consistent with TG utilization during metamorphosis (**SFigure 3D**). Furthermore, whole animal glycogen was even lower in 7-10 days old male and female *FASN1^FB-RNAi^* adult flies than in controls. This reduction may reflect utilization of residual glycogen stores to sustain early post-eclosion *FASN1^FB-RNAi^* adults, in the absence of LDs (**SFigure 3E**). Taken together, these findings reveal a dramatic shift from TG to glycogen storage in the FB of *FASN1^FB-RNAi^*larvae and imply the functional importance of glycogen synthesis and mobilization for successful completion of development.

### A cell autonomous TG:glycogen metabolic switch is triggered specifically by DNL blockage and rescued by palmitate

Elevated glycogen storage and utilization when TG stores were limited indicated a TG:glycogen metabolic switch consistent with rewiring of metabolism described in other contexts^58,59^. To understand if this TG:glycogen switch was cell autonomous, we utilized the ubiquitously expressed mosaic driver *da-Gal4* that activates a *UAS-RNAi* transgene in a subset of FB cells^60^. Indeed, *da-Gal4>FASN1^RNAi^* larval FBs exhibited a mosaic pattern of LD-depleted cells adjacent to lipid engorged cells (**SFigure 4A**). In line with this, PAS stained *da-Gal4>FASN1^RNAi^* FBs exhibited a similar mosaic pattern of glycogen accumulation, with dark PAS staining confined to *FASN1*-knockdown cells (**SFigure 4B, bottom panel, white arrows**) while neighboring unaffected cells resembled those of controls (**SFigure 5B, light blue arrows**). This indicated the TG:glycogen metabolic switch was a cell autonomous effect of early DNL blockage.

Since glycogen elevation was observed following the dampening of an early step in DNL (i.e. FASN1 depletion), we examined whether perturbing other steps in the DNL pathway similarly affected glycogen accumulation in the FB. We took a candidate-based approach and RNAi depleted several enzymes across the DNL pathway. Of note, RNAi-mediated depletion of acetyl-CoA carboxylase (ACC), the rate-limiting enzyme of DNL acting immediately upstream of FASN1, resulted in visually pronounced LD depletion and a dramatic increase in glycogen-associated PAS staining intensity, akin to that observed in *FASN1^FB-RNA^*^i^ FB (**SFigure 4C,D**). In contrast, FB-specific depletion of enzymes acting downstream of FASN1, including acyl-CoA ligase ACSL, the acyltransferase GPAT4, dLipin, and the TG synthesis enzyme Midway (Mdy), did not result in a substantially darker PAS signal (**SFigure 4D**). In line with this, these RNAi lies also led to a partial but milder LD depletion (**SFigure 4C**). Based on these observations, the apparent correlation between the extent of LD loss and enhanced glycogen staining suggests that the balance between lipid and glycogen stores may be influenced by fatty acid biosynthesis and TG storage.

ACC and FASN1 catalyze the first two steps in DNL, generating malonyl-CoA and palmitate (subsequently converted to palmitoyl-CoA), respectively, whereas downstream enzymes modify these fatty acyl-CoAs into lyso-lipids and mature lipids for the cell^34,61^ (**SFigure 1A,4C**). We hypothesized that *fatty acids themselves* may regulate the TG:glycogen balance in the FB, given that ACC or FASN1 RNAi depletion generated such potent glycogen accumulation. Since palmitate is the major enzymatic product of FASN1^62^, we supplemented *Drosophila* food with 10% palm oil (PO), which contains approximately ∼44% palmitate^63^, to test whether dietary palmitate could suppress the TG-to-glycogen metabolic switch in *FASN1^FB-RNAi^* animals. Significantly, *FASN1^FB-RNAi^*larvae reared on food enriched with 10% PO exhibited substantial restoration of LDs and TG stores (**Figure 4A,B**), and a concomitant reduction in glycogen to those comparable to controls (**Figure 4C,D**). These findings suggest that PO was sufficient to attenuate the shift in energy storage induced by FASN1 knockdown, favoring lipid storage over glycogen accumulation.

**Figure 4:**
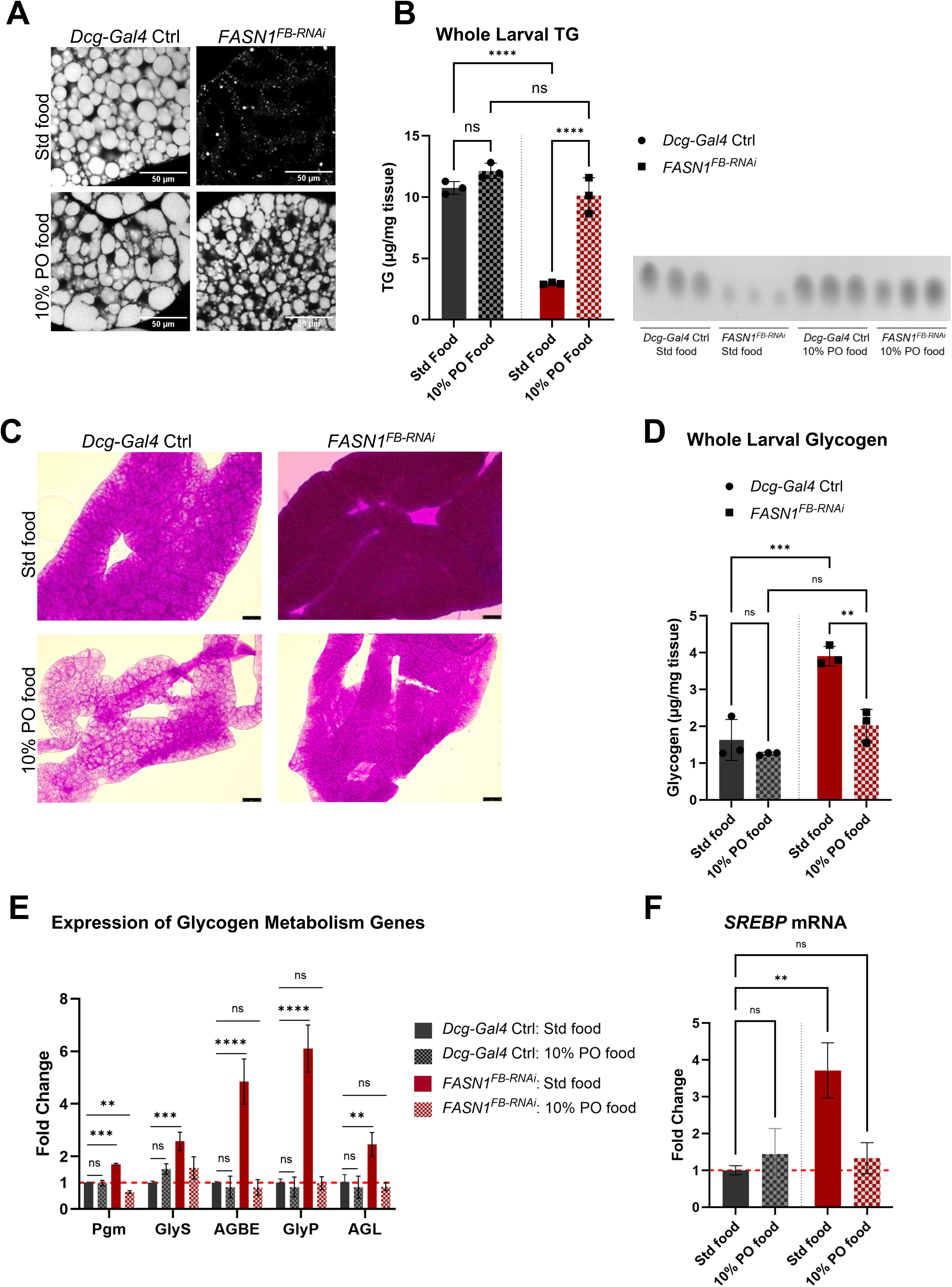
Disruption of fatty acid synthesis during early *de novo* lipogenesis (DNL) drives a metabolic rewiring that favors glycogen storage over triglyceride accumulation. A) Confocal images of MDH-stained LDs in FBs from *Dcg-Gal4* control and *FASN1^FB-RNAi^* L3 larvae cultured on standard food and food supplemented with 10% Palm Oil (PO). Scale bar 50 µm. B) Total triglycerides (TG) measured by thin layer chromatography (TLC) in *Dcg-Gal4* control and *FASN1^FB-RNAi^* L3 larvae cultured on standard and 10% PO food. TLC plate representing TG bands corresponding to the quantified samples is shown alongside the graph (sample volumes were normalized to larval weight). Data are shown as mean ± SD from 3 biological replicates per genotype (15 larvae per replicate). P values were determined using Welch’s one-way ANOVA. C) PAS glycogen staining in FBs from *Dcg-Gal4* control and *FASN1^FB-RNAi^* L3 larvae cultured in standard and 10% PO food. Scale bar 50 µm. D) Biochemical quantification of glycogen extracted from *Dcg-Gal4* control and *FASN1^FB-RNAi^* whole L3 larvae reared on standard and 10% PO food. Data are shown as mean ± SD from 3 biological replicates per genotype (15 larvae per replicate), normalized to larval weight. P values were determined using two-way ANOVA followed by Šídák’s multiple-comparisons test. E) qPCR analysis of glycogen metabolism gene expression in FBs of *Dcg-Gal4* control and *FASN1^FB-RNAi^* L3 larvae reared on standard and 10% PO supplemented-food. Expression levels are shown relative to those in *Dcg-Gal4* controls on standard food. Data are presented as mean ± SD from 3 biological replicates per genotype (FBs pooled from 20-30 L3 larvae per replicate). P values were determined using an unpaired, two-tailed Student’s t-test. F) Relative *SREBP* mRNA expression in FBs of *Dcg-Gal4* control and *FASN1^FB-RNAi^* L3 larvae raised on standard and 10% PO food. Expression levels are normalized to those in *Dcg-Gal4* controls on standard food. Data are presented as mean ± SD from 3 biological replicates per genotype (FBs pooled from 20-30 L3 larvae per replicate). P values were determined using an unpaired, two-tailed Student’s t-test.

Since our proteomic analysis and qPCR indicated upregulation of enzymes involved in both glycogenesis and glycogenolysis in FASN1-deficient FBs, we also queried whether larvae fed on PO supplemented food displayed altered carbohydrate-associated mRNA transcripts. Consistent with the reduced glycogen staining, qPCR analyses of FBs from *FASN1^FB-RNAi^* larvae revealed that animals fed 10% PO food had suppressed expression of glycogen related genes such as *GlyS* or the glycogen debranching enzyme *AGL* compared to those reared on normal food (**Figure 4E)**. This further indicated that dietary fatty acids were sufficient to modulate glycogen metabolism in the FB, suggesting a trade-off between lipid and glycogen storage.

Since dietary fatty acids could influence transcriptional remodeling in FASN1-depleted FBs, we considered what transcription factors could regulate this response. Previous work on the *Drosophila* transcription factor Sterol Regulatory Element-Binding Protein (SREBP) showed that palmitate directly inhibited SREBP proteolytic processing and transcriptional activation^64^. According to the canonical model, palmitate incorporation into membrane phospholipids, including phosphatidylethanolamine (PE), influenced ER membrane fluidity and inhibited SREBP release from the ER and subsequent cleavage at the Golgi, required for its nuclear translocation and transcriptional activity^64,65^. Given that *FASN1* KD impairs *de novo* fatty acid synthesis, and that proteomic analysis revealed increased SREBP protein in FASN1-depleted fat bodies (**Figure 3B**, **SFigure 4E**), we next performed qPCR in FBs from control and *FASN1^FB-RNAi^* FB larvae reared on standard and 10% PO food, to assess whether dietary fatty acid supplementation modulated *SREBP* mRNA levels. Indeed, *SREBP* transcripts were significantly elevated in *FASN1^FB-RNAi^* FB under standard dietary conditions but reverted to control levels in larvae reared on food supplemented with 10% PO (**Figure 4F**). These data suggest that altered lipid availability may influence SREBP regulation, hinting at a role for SREBP in glycogen storage under conditions of impaired fatty acid synthesis.

### *Drosophila* SREBP is a mediator of the TG:glycogen metabolic switch

Whereas mammalian SREBPs regulate fatty acid (SREBP-1) and cholesterol (SREBP-2) biosynthesis, the single *Drosophila* SREBP homolog canonically regulates only fatty acid synthesis^64–66^. However, beyond their roles in lipid metabolism, SREBP homologs have also been implicated in glucose homeostasis^41–44^. Notably, previous literature supports a poorly understood potential link between SREBP and carbohydrate storage in the mammalian liver. For example, mammalian SREBP-1-deficient hepatocytes display lower glycogen levels in the fed state, suggesting a role for SREBP-1 in liver carbohydrate storage^67^. We therefore hypothesized that *Drosophila* SREBP may be required for the TG:glycogen switch observed when DNL is perturbed in the FB. To test this, we first investigated whether SREBP was required for the boosted glycogen levels observed in FASN1-depleted FBs. Glycogen accumulation was evaluated in larval FBs from control animals, *FASN1^FB-RNAi^* or *SREBP^FB-RNAi^*single-KDs, or the double KD of *FASN1^FB-RNAi^* and *SREBP^FB-RNA^*, using both PAS glycogen staining and biochemical measurements (**Figure 5A,B**). Importantly, KD efficiency was validated by qPCR for each RNAi line **(SFigure 5A**). As expected, glycogen levels were elevated in the FB of *FASN1*-depleted larvae. However, strikingly simultaneous loss of *SREBP* and *FASN1* reduced FB glycogen stores by over 80% relative to *FASN1^FB-RNAi^* alone, and by∼50% relative to controls (**Figure 5A,B**). Of note, this glycogen reduction occurred despite similarly low LD and TG abundance in *FASN1^FB-RNAi^* single KDs and *SREBP^FB-RNAi^ + FASN1^FB-RNAi^* double KDs, suggesting the “rescue” of glycogen levels was not simply due to a restoration of FB lipid stores (**SFigure 5B,C**). Furthermore, qPCR analysis of FBs underscored our hypothesis that the upregulation of genes involved in glycogenesis and glycogenolysis in *FASN1^FB-RNAi^* FBs may require SREBP. For example, mRNA transcripts of key glycogen-associated factors including GlyS, AGBE, GlyP, and AGL were significantly elevated in *FASN1^FB-RNAi^*FBs, but these returned to near baseline levels in *SREBP^FB-RNAi^ + FASN1^FB-RNAi^* samples (**Figure 5C**). Additionally, expression of the putative alpha-glucosidase *tobi*, one of the highly induced genes in *FASN1^FB-RNAi^* FBs, was reduced by ∼75% in *SREBP^FB-RNAi^ + FASN1^FB-RNAi^* KDs (**SFigure 5D**). Collectively, this indicated that *Drosophila* SREBP was required for the boost in glycogen storage in the absence of FASN1 in the FB.

**Figure 5:**
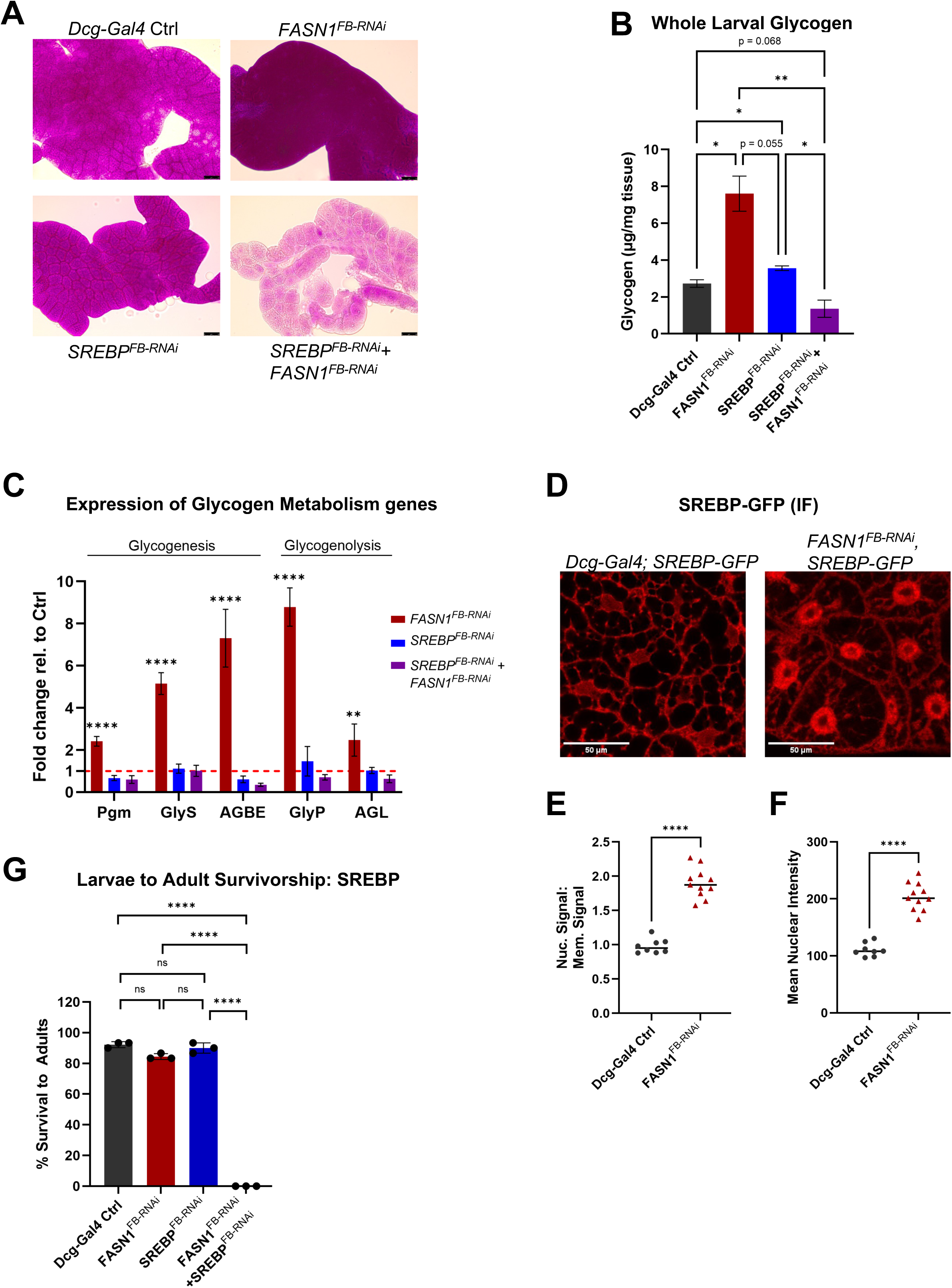
*Drosophila* SREBP is a critical determinant of the TG-to-glycogen storage shift. A) PAS staining of glycogen in FBs of L3 larvae following loss of function of *FASN1*, *SREBP*, or both, alongside *Dcg-Gal4* controls. Scale Bar 50 µm. B) Biochemical quantification of glycogen extracted from *Dcg-Gal4* control, *FASN1^FB-RNAi^* and *SREBP^FB-RNAi^* single and *FASN1^FB-RNAi^*+*SREBP^FB-RNAi^* double KD whole L3 larvae. Data are shown as mean ± SD from 3 biological replicates per genotype (15 larvae per replicate), normalized to larval weight. P values were determined using Welch’s one-way ANOVA. C) Relative expression of glycogen metabolism genes in FBs of *Dcg-Gal4* control, *FASN1^FB-RNAi^*, *SREBP^FB-RNAi^*, and *FASN1^FB-RNAi^* +*SREBP^FB-RNAi^* L3 larvae. Data are shown as mean ± SD from 3 biological replicates per genotype (FBs pooled from 20-30 L3 larvae per replicate). P values were determined using an unpaired, two-tailed Student’s t-test. D) Confocal imaging of the active N-terminus SREBP-GFP fragment in *Dcg-Gal4* control and *FASN1^FB-RNAi^* FB cells, detected by immunofluorescence (IF) staining using an anti-GFP primary antibody and an RFP-conjugated secondary antibody. Scale bar 50 µm. E) Mean nuclear SREBP-GFP IF intensity in *Dcg-Gal4* control and *FASN1^FB-RNAi^* FB cells, quantified using Fiji ImageJ. Data points represent mean ± SD from 8 *Dcg-Gal4* control and 11 *FASN1^FB-RNAi^* FB tissues (24 cells per genotype). P values were determined using an unpaired, two-tailed Welch’s t-test. F) Ratio of nuclear to membrane-associated SREBP-GFP IF signal in *Dcg-Gal4* control and *FASN1^FB-RNAi^* FBs, quantified using Fiji ImageJ. Data points represent mean ± SD from 8 *Dcg-Gal4* control and 11 *FASN1^FB-RNAi^* FB tissues (24 cells per genotype). P values were determined using an unpaired, two-tailed Welch’s t-test. G) Percentage of L3 larvae (*Dcg-Gal4* control, *FASN1^FB-RNAi^*, *SREBP^FB-RNAi^*, *SREBP^FB-RNA^ ^i^*+ *FASN1^FB-RNAi^*) successfully completing development to yield adult flies. Data are shown as mean ± SD from 3 independent experiments (90 L3 larvae per genotype across 3 experimental repeats). P values were determined using a two-tailed Fisher’s exact test on pooled eclosion counts.

To more directly investigate whether SREBP was activated in FASN1-depleted FB cells, we directly monitored SREBP subcellular localization. Inactive SREBP is localized to the ER membrane, but upon activation, the SREBP–SCAP complex translocates to the Golgi, where proteolytic cleavage releases the N-terminal transcription factor, which subsequently enters the nucleus^68^. To more directly examine whether SREBP activation was increased in *FASN1^FB-RNAi^* FB cells, we crossed control and *FASN1^FB-RNAi^* flies with a transgenic line expressing *Drosophila* SREBP tagged with GFP at its N-terminus, enabling visualization of the nuclear localization of the cleaved, transcriptionally active N-terminal fragment. We performed immunofluorescence staining using an anti-GFP primary antibody and an RFP-conjugated secondary antibody to detect SREBP-GFP. As expected, confocal imaging revealed primarily ER-localized SREBP-GFP in control FBs, with some detectable nuclear SREBP-GFP, reflecting baseline SREBP activity (**Figure 5D**). Strikingly, there was significantly greater nuclear SREBP-GFP signal in *FASN1^FB-RNAi^*FB cells than in control FB cells (**Figure 5D,E,F**), consistent with increased SREBP activation upon depletion of DNL. This supported the model where loss of FASN1-derived fatty acids promotes SREBP activation and subsequent glycogen storage, suggesting a noncanonical role for *Drosophila* SREBP in FB glycogen metabolism.

From this we hypothesized that SREBP would be essential for adult *Drosophila* survival in the absence of normal FB FASN1 activity. We investigated whether SREBP was required for *Drosophila* development in *FASN1^FB-RNAi^*animals by depleting *SREBP* either alone or in combination with *FASN1* in the FB. Strikingly, whereas SREBP loss alone did not impact *Drosophila* development into flies, *FASN1* and *SREBP* dual RNAi depletion led to a complete failure of pupation with no viable adults (**Figure 5G**). These findings suggest that a noncanonical, SREBP-dependent metabolic switch promotes glycogen accumulation in response to impaired fatty acid synthesis, potentially supporting organismal survival during *Drosophila* development.

## Discussion

Energy balance is largely reliant on glucose availability and mobilization, which drives adipose tissue metabolism and storage of substrates as TG and glycogen^3,4,69^. Dietary sugars stimulate expression of *de novo* lipogenic enzymes including ACC and FASN that convert acetyl CoA derived from pyruvate into fatty acids that can be esterified to TG and assimilated into LDs^5,18^. In *Drosophila,* the FB is the central lipid depot, and performs liver-like functions to store both TG and glycogen^18,69^. We capitalized on this TG:glycogen dual availability to identify factors mediating energy balance between them, and adaptation in animal development. We find that FB-specific depletion of FASN1 leads to a near-complete loss of TG and re-orienting towards carbohydrate-dependent metabolism. Here we provide evidence for a cell autonomous TG-to-glycogen metabolic switch in *Drosophila* triggered by a genetic disruption of early DNL fatty acid synthesis. This shift is characterized by a dramatic increase in FB glycogen accumulation and utilization, which together ensures development in the absence of abundant fat stores. Using genetics and biochemical assays, we show this switch requires *Drosophila* SREBP, a lipid sensing and nutrient regulating transcription factor that traditionally regulates lipogenesis but has been observed to also influence carbohydrate metabolism in the mammalian liver. Critically, in *Drosophila* palmitate inhibits SREBP proteolytic processing required for its activation. In line with this, our data suggest a *feedback loop* where reduced fatty acid availability upon DNL blockage activates SREBP to promote metabolic rewiring that boosts glycogen metabolism for organismal survival.

*Drosophila* must reach critical weight to achieve pupation and metamorphosis, and the biomass for this has been proposed to come from FB lipids^20–22^. Our findings challenge this notion as we find *FASN1^FB-RNAi^* larvae lack lipid stores yet grow to full size and pupate without developmental delay. GO term analysis performed on proteomics data indicated significant reprogramming of glucose metabolism, whereby nutrients were channeled back into carbohydrates via glycogenesis We hypothesize that the resulting 20-fold increase in glycogen deposits in *FASN1^FB-RNAi^*FBs may contribute to increased body mass by promoting water retention within FB cells. Indeed, glycogen absorbs 3-5 times its weight in water which may help compensate for lipid loss and help *FASN1^FB-RNAi^*larvae to reach critical weight and size^70^. Additionally, proteomics and transcriptomics identified tobi, an alpha-glucosidase, as a potential contributor to increased carbohydrate utilization in response to TG depletion in FB tissue. Our data indicate that glycogen reserves, generally labelled a ‘metabolic safeguard’ and deemed dispensable for metamorphosis^6,8^, are essential for *FASN1^FB-RNAi^*animals during pupal to pharate adult transition, while control animals rely mostly on TG. It is known that FB glycogen, can be mobilized to replenish trehalose in circulation during nutrient shortages^71^, suggesting that both FB glycogen and hemolymph trehalose support fly development in the absence of TG. Indeed, we detected higher amounts trehalose in *FASN1^FB-RNAi^* larval hemolymph, probably owing to enhanced glycogen breakdown in these animals. This metabolic rewiring reveals a previously unappreciated versatility to *Drosophila* development.

The absence of apparent growth defects in *FASN1^FB-RNAi^*larvae, together with their timely metamorphosis into full-sized adults, implied that developmental growth programs remain largely intact despite profound lipid depletion. Consistent with this, *FASN1^FB-RNAi^* larvae maintained normal circulating glucose levels, the hemolymph carbohydrate fraction typically more sensitive to metabolic cues^16^ and exhibited no apparent defects in food intake. Interestingly, palmitate is the major fatty acid produced by FASN1^62^. Therefore, we hypothesized that DNL inhibition lowers palmitate in the FB, thereby impairing TG synthesis and possibly triggering a compensatory metabolic switch towards glycogen synthesis. In support of this, culturing *FASN1^FB-RNAi^* larvae on a palmitate-rich diet restored FB TG stores and suppressed glycogen accumulation. Fatty acid availability is a relevant regulatory feature of *Drosophila* SREBP, which responds to intracellular changes in palmitate and related membrane lipids^64,65^. We therefore asked whether SREBP might mediate the metabolic response to DNL inhibition.

Mammals possess three SREBPs, isoforms SREBP-1a andSREBP-1c, and related homolog SREBP-2. SREBP-1a and SREBP-1c support fatty acid and TG synthesis through insulin-dependent lipogenesis, whereas SREBP-2 primarily regulates membrane sterol homeostasis^72–74^. In contrast, *Drosophila* encodes a single SREBP homolog that recapitulates many functions of its mammalian counterparts, transcriptionally activating DNL genes, notably ACC and FASN1, under feeding conditions to promote palmitate synthesis and feedback regulation of SREBP activity^64,65,75^. Although carbohydrate metabolism is not considered a canonical function of SREBP, studies in mice have implicated these transcription factors in hepatic glucose homeostasis. These findings prompted us to investigate whether *Drosophila* SREBP has a potentially conserved role in glucose regulation, particularly in the control of glycogen storage. We hypothesized that DNL inhibition lowers palmitate levels in the FB, thereby promoting SREBP activation. Consistent with this possibility, both SREBP mRNA and protein levels were elevated in *FASN1^FB-RNAi^*FBs. Notably, SREBP single KD did not reduce glycogen levels and instead showed a mild increase relative to controls, suggesting a context-dependent role for SREBP in glycogen regulation. However, concomitant *SREBP* RNAi-depletion in the *FASN1^FB-RNAi^* background drastically reduced glycogen accumulation compared with *FASN1^FB-RNAi^* single KD FBs, suggesting that SREBP is required for the shift toward glycogen storage in response to TG depletion. Notably, this reduction occurred despite persistent TG deficiency in *SREBP^FB-RNAi^* + *FASN1^FB-RNAi^* double KD FBs.

We further hypothesized that SREBP-mediated metabolic rewiring in palmitate-deficient *FASN1^FB-RNAi^* FBs involves the transcriptional activity of SREBP. Upon activation, SREBP translocates to the nucleus, where it may regulate the expression of genes involved in glycogen synthesis and breakdown. Indeed, the expression of genes regulating glycogen metabolism, which was elevated in *FASN1^FB-RNAi^*FBs, were restored to control levels in this background following either SREBP depletion or dietary supplementation with palm oil. Further supporting enhanced SREBP activity and metabolic rewiring in response to DNL inhibition, we observed significantly greater nuclear localization of the GFP-tagged active N-terminal SREBP fragment in *FASN1^FB-RNAi^* FB cells compared with controls. We propose that in the FB *Drosophila* SREBP lipid-sensing capabilities may play a more important role in regulating glycogen metabolism (**Figure 6**). A few studies in mice have reinforced a noncanonical role for SREBP in regulating glycogen metabolism. Intriguingly, overexpression of the active form of SREBP-1c in the liver of diabetic mice increased hepatic glycogen stores, associated with SREBP-1c-driven transcriptional changes in glucose metabolism genes^76^. Similarly, SREBP1 promoted glycogen accumulation in obese mice through upregulation of the glycogenic scaffolding protein PTG^76,77^. In contrast, Ruiz *et al.* (2013) reported that silencing SREBP-1 in the mouse liver reduced glycogen deposits and blunted the expression and activity of glycogenic factors^78^.

**Figure 6:**
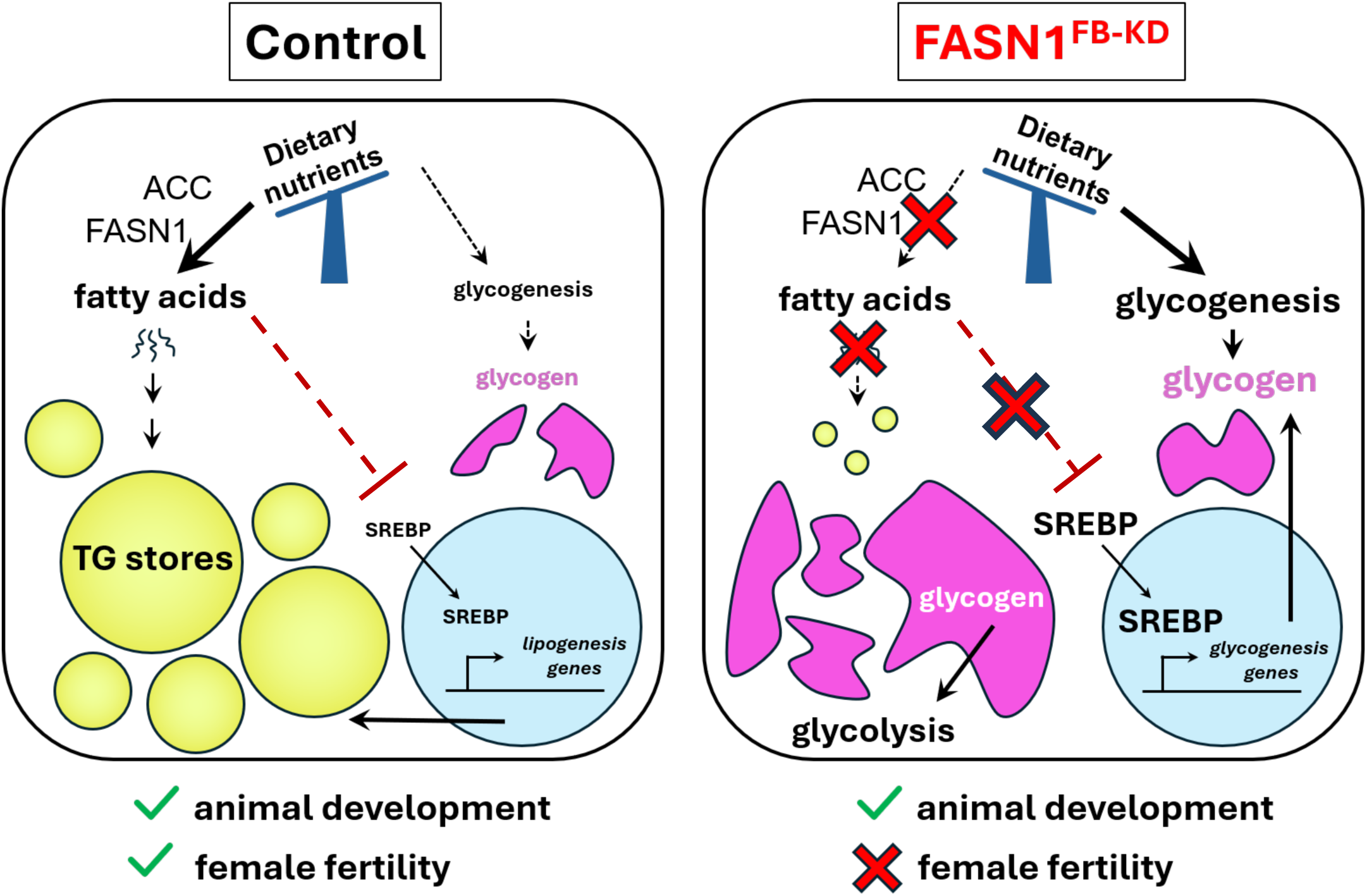
Working model for how triglyceride:carbohydrate cross-talk and metabolic re-wiring occurs in a ‘fat-less’ *Drosophila* model to promote growth and development

Interestingly, FASN1 depletion was also associated with increased expression of glycolytic genes, including Pfk and Pyk. Increased glycogen mobilization in these FBs likely results in elevated free glucose availability, which can be channeled into glycolysis to support catabolism and ATP generation. Moreover, previous studies in mammals have implicated SREBP-1 in the regulation of glycolytic gene expression, including glucokinase (GK) and Pyk^41,64,79^. These observations raise the possibility that increased SREBP activity contributes to increased glycolysis as part of the broader transcriptional remodeling of carbohydrate metabolism that increases glycogen accumulation in lipid depleted *FASN1^FB-RNAi^*FBs. In support of this, mammalian ATPCL (ACLY), which we find boosted in FASN1-depleted FBs, is an established transcriptional target of SREBP proteins^80,81^, and generates cytosolic acetyl-CoA, an essential precursor of de novo fatty acid synthesis. However, a study by Seegmiller et al. did not detect altered ATPCL expression in *Drosophila* S2 cells^64^. Thus, the marked increase in ATPCL expression observed in FASN1-depleted FBs may represent a compensatory response to impaired fatty acid synthesis, potentially aimed at maintaining cytosolic acetyl-CoA availability in lipid depleted tissue. Whether these changes in glycolytic gene expression and ATPCL levels contribute functionally to the TG-to-glycogen shift in FASN1-deficient animals remains to be determined.

Fat-depleted *Drosophila* maintain developmental integrity through rebalancing fat:carbohydrate metabolism, but does this expose new vulnerabilities? *FASN1^FB-RNAi^* male and female flies exhibited barely detectable TG stores and half the glycogen reserves of controls; unsurprisingly this rendered them starvation sensitive and both males and females displayed shorter median lifespans relative to controls. Fertility was adversely affected in *FASN1^FB-RNAi^* females. *FASN1^FB-RNA^*ovaries appeared morphologically underdeveloped, which may be a consequence of a phenomenon called “organ sparing”. When energy supply is limited, animals preferentially support development of certain organs at the cost of others^50^. In *FASN1^FB-RNAi^* female flies, we noted that egg chamber oocyte selection from nurse cells and it’s subsequent wrapping by follicle cells, both occurred improperly, arresting oogenesis at developmental stage 9-10. Since maternal TG deposition is a critical checkpoint for oocyte maturation and embryo survival, failure of oogenesis in *FASN1^FB-RNAi^* females likely results from low level of LDs in nurse cells. TG stored in LDs of nurse cells is synthesized from DAG, 80% of which is exogenously resourced from free-floating cells of the dissociated larval FB that persist in the newly hatched fly^24,54^. We reason that lipophorin-mediated DAG delivery from *FASN1^FB-RNAi^* lipid-deficient fat cells to the ovaries was severely limited. In line with this, subsequent apoptosis of egg chambers due to low lipid content has been reported by other studies^82,83^.

In closing, the existence of a TG:glycogen metabolic switch, regulated by a SREBP-dependent feedback loop, in the *Drosophila* FB suggests versatile adaptive capacity for the *Drosophila* bioenergetics developmental program. Similar TG:glycogen rewiring is anecdotally also observed in mammals. In mice when TG storage was genetically blocked in adipose by DGAT1 and DGAT2 ablation, mice exhibit modest but significant increases in glycogen storage^84^. Furthermore, human organs exhibit specific ratios of fat and glycogen nutrient reserves, but how these are balanced and adapt to changes in nutrient availability remains incompletely understood. Here, we provide evidence for unique metabolic rewiring in the fruit fly which may apply to diverse human tissues. It suggests remarkable plasticity to nutrient storage and utilization in the struggle to adapt and survive.

## Acknowledgements

We thank members of the Henne and DeBerardinis labs for helpful advice and discussions during the development of this study. We also thank Anastasiia Kovalenko for initial help profiling fat body LD phenotypes. This work is supported by grants from the Welch Foundation I-1873 (W.M.H), the National Institutes of Health (NIH) grants GM119768 (W.M.H), DK126887 (W.M.H.), the Ara Parseghian Medical Research Fund, and the UT Southwestern Endowed Scholars Program. T.P.M. and R.J.D. are supported by CPRIT grant RP180778. R.J.D. is supported by NIH grants R35CA220449 and P50CA196516. L.G.Z., T.P.M., and the CRI Metabolomics Facility are supported by the Cancer Prevention Research Institute of Texas (CPRIT Core Facilities Support Award RP240494).

## Data and Information Availability

All data related to this study is available in the materials and also available upon request.

## MATERIALS AND METHODS

### Fly cultures and strains

All *Drosophila* animal research was performed in accordance with NIH recommended policies. *Drosophila* late-feeding third instar (L3) larvae (∼100-108 h After Egg Laying or AEL) were used for fat tissue microscopy and biochemical assays, unless otherwise stated. For lifespan and starvation assays, adult male and female flies were used. Flies were reared and maintained on standard fly food containing cornmeal, yeast, molasses, and agar. The *Dcg-Gal4* fat body (FB)-specific driver^46^ was provided by Jonathan M. Graff (former UTSW). *UAS-FASN1^RNAi^* and all other *TRiP RNAi* lines, protein/organelle marker strains and other transgenic stocks used in this study were obtained from the Bloomington Stock Center (Bloomington, IN). Typically, 5 to 7 *Gal4* driver males were crossed with ∼20-25 females from test *UAS* or control lines *(UAS-mCherry^RNAi^)*. All experimental crosses and controls were cultured at ∼100-150 animals per vial (mid-density, which maintains metabolic stability^85^), in a fly incubator with a 12-hour light/dark cycle and maintained at 25°C for uniform transgene expression. Please see Table S1 for a list of *Drosophila* lines used.

### Larval fat body staining

Late feeding L3 larvae were gently removed from inside the food media using a paintbrush and rinsed in water to remove food particles. Larval fat bodies (FBs) or guts were dissected in PBS using Dumont #5 forceps (Electron Microscopy Sciences #0103-5-TF). All tissues were fixed in 4% paraformaldehyde (PFA) for 20 min at room temperature and rinsed briefly in PBS prior to staining. Lipid droplets in FBs were stained by incubating FBs in 100 µM monodansylpentane (MDH) LD stain (Absepta #SM1000a) for 20 min at room temperature, in the dark. Next, FBs were rinsed in PBS and mounted on slides in SlowFade Gold antifade reagent with nuclear DAPI (Invitrogen #S36936) for subsequent imaging.

### Female ovary dissection and staining

Ovaries from 7 to 10-day old adult females were dissected in PBS, fixed in 4% PFA for 20 min at room temperature. Following a brief wash in PBS, ovaries were incubated in CellMask Actin stain (1:1000, Invitrogen #A57243) or Phalloidin (1:400 in PBS-T, Invitrogen #A30104) for staining of F-actin structures, according to manufacturer’s instructions. LDs in ovaries were stained with BODIPY 493/503 (1:500, Invitrogen #D3922) for 30 min in the dark. Excess stain from ovaries was removed by washing in PBS, then mounted in SlowFade Gold antifade reagent containing DAPI for imaging. Egg chambers within ovarioles of the ovaries were staged according to established morphological criteria described in literature^86–88^.

### Confocal fluorescence microscopy

Prepared slides were imaged using Zeiss LSM880 inverted laser scanning confocal microscope. Tissues stained with organelle dyes (LD: MDH/ BODIPY; F-actin: CellMask, Phalloidin), RFP-conjugated secondary antibody and those expressing transgenic fluorescent-tagged proteins (e.g., UAS-GFP; ER marker *UAS-Sec61B-TdT*; PM markers *UAS-mcd8-GFP* and *UAS-Glut4-GFP*) were imaged using the appropriate channel filter DAPI/GFP/RFP. Most FBs were imaged using a 40X oil immersion objective, and adult female ovaries with 10X or 20X objective.

### Light and other fluorescence microscopy

Whole L3 larvae without or with GFP-expressing (*Dcg-Gal4>UAS-GFP*) FBs were imaged using the Leica S8AP0 stereoscope and EVOS FL Cell Imaging System (Thermo Fisher), respectively. Adult male and female flies, and pupae were also imaged on Leica S8AP0 stereoscope.

### Transmission electron microscopy

FB tissue was removed from *Drosophila* larvae, fixed in 2.5% (v/v) glutaraldehyde in 0.1 M sodium cacodylate buffer paraformaldehyde, and processed in the UT Southwestern Electron Microscopy Core Facility. They were post-fixed in 1% osmium tetroxide and 0.8% K3[Fe(CN6)] in 0.1 M sodium cacodylate buffer for 1 h at room temperature. Cells were rinsed with water and en bloc stained with 2% aqueous uranyl acetate overnight. Next, they were rinsed in buffer and dehydrated with increasing concentration of ethanol, infiltrated with Embed-812 resin and polymerized in a 60°C oven overnight. Blocks were sectioned with a diamond knife (Diatome) on a Leica Ultracut UCT (7) ultramicrotome (Leica Microsystems) and collected onto copper grids, post stained with 2% aqueous uranyl acetate and lead citrate. Images were acquired on a JOEL 1400 Plus transmission electron microscope using a voltage of 120 kV.

### Quantitative PCR

Whole RNA was extracted from FB tissues (20-30 larvae per biological replicate, n = 3) using Trizol reagent (Invitrogen #15596026) according to manufacturer’s instructions. RNA was quantified by Denovix spectrophotometer, and 1 µg of RNA was reverse transcribed to cDNA using iScript Reverse Transcriptase Supermix for RT-qPCR Kit (BioRad #1708841). qPCR was performed with cDNA using the SsoAdvanced Universal SYBR Green Supermix (BioRad #1725274) on the BioRad CFX96 Real-Time System; mRNA expression data were normalized to that of the fly housekeeping gene rp49. Primer sequences for all transcripts amplified are available upon request.

### Glucose and Trehalose assays

Feeding larvae washed out from culture vials were collected in 630□µm (Genesee #NC9432449) mesh-fitted baskets (Genesee #46-102) and rinsed to get rid of adherent food particles. Larvae were dried and divided onto into piles (10–12 larvae each, n = 3) on a strip of parafilm. Larvae were bled by tearing the cuticle with Dumont 5 forceps (Electron Microscopy Sciences). Two µl of colorless hemolymph was aspirated from each pile and separately transferred to 96-well plates (Thermo Scientific #14-245-71) containing 0.1% N-Phenylthiourea (Sigma-Aldrich #P7629) in 50□µl PBS. 150□µl of Autokit Glucose reagent (Fujifilm #994-03011) was added to each well and incubated at room temperature for 20□min before measuring absorbance at 505 nm. Glucose concentration was calculated from a standard curve generated with manufacturer’s glucose standards. For trehalose assays, 8□µl of dilute hemolymph was treated with 5□µl of (diluted 8X) porcine kidney trehalase (Sigma-Aldrich #T8778) overnight at 37□°C. Ten µl of treated sample was assayed for trehalose as described for glucose. Trehalose amounts were calculated from standard glucose, as for glucose.

### Glycogen assay

Whole animals (10-15 larvae/pharate pupae/adults per biological replicate, n = 3) or dissected FBs (25-30 larvae per biological replicate, n = 3) were homogenized in 300 µl ice-cold PBS using a pestle or sonication and syringe (29G1/2 needle), respectively. After reserving 20 µl of the homogenate for protein Bradford quantification, the rest of the homogenate was heat inactivated at 70°C for 10 min. Homogenate was then centrifuged at maximum speed at 4°C for 10 min and supernatant was collected in a new tube. In a 96-well plate, 30 µl of each sample was loaded in duplicate rows. Then, 100 µl of Autokit Glucose reagent + amyloglucosidase (2 µl amyloglucosidase {Sigma #A1602; 25mg} per 1 ml of Glucose reagent) was added to one row of samples, and 100 µl of Glucose reagent alone (without amyloglucosidase) was added to the duplicate row of samples, and to the glucose standards. The plate was incubated at 37°C for 30 min, after which color intensity was measured using a microplate reader at 505 nm. Free glucose concentration in treated and untreated samples was calculated based on the glucose standard curve. Glycogen concentration (as free glucose) was determined by subtracting glucose in the untreated samples from those treated with amyloglucosidase. Finally, total glycogen was normalized to protein (for FBs) measured by the Bradford assay using Bio-Rad Protein Assay Dye Reagent Concentrate (#5000006) or normalized to weight (for whole animals).

### Periodic Acid-Schiff stain

Glycogen in FB tissues from late feeding L3 larvae was stained using the Periodic Acid-Schiff (PAS) kit (Sigma Diagnostics #SD395B). Larval FBs were dissected in 1% BSA in PBS, fixed with 4% paraformaldehyde for 20 min, and washed twice with 1% BSA in PBS. FBs were then incubated in 0.5% period acid solution for 5 min, washed twice with 1% BSA in PBS, then stained with Schiff’s reagent for 15 min, washed again, and mounted on slides. PAS-stained FB tissues were imaged using the Leica DM6B light microscope. Comparative PAS staining intensity was quantified using Fiji ImageJ software.

#### Food Ingestion Assay

Larvae cultured on Coomassie dye-supplemented food (200 µg dye/ml media) were harvested, washed in water, blotted dry, and transferred to 1.5□ml Eppendorf tubes on ice (30 per biological replicate, n = 3). Larvae were homogenized in 200□µl of 100% methanol. Samples were centrifuged at 14,000□rpm at 4°C for 10□min; 175□µl of the supernatant was transferred to fresh tubes containing 175□µl of ddH₂O and centrifuged again. Absorbance of this supernatant was measured at 595□nm, and dye amount (food ingested) was calculated by comparing with a standard curve generated from a range of Coomassie dye concentrations.

### Thin layer chromatography

Late feeding L3 larvae or adult flies (10-15 per biological replicate, n = 3) were weighed, then homogenized in 2:1:0.8 of methanol:chloroform:water. Samples were incubated in a 37°C water bath for 1 h. Chloroform and 1M KCl (1:1) were added to the sample, centrifuged at 3000 rpm for 2 min, and the bottom layer containing lipids was aspirated using a syringe. Lipids were dried using argon gas and resuspended in chloroform (100 µl of chloroform/7 mg of fly weight). Extracted lipids alongside serially diluted standard neutral lipids of known concentrations were separated on TLC plates using hexane:diethyl ether:acetic acid solvent (80:20:1, v/v/v). TLC plate was air dried for 10 min, spray stained with 3% copper (II) acetate in 8% phosphoric acid and incubated at 145°C in the oven for 30 min to 1 h to allow bands to develop for scanning and imaging. Neutral lipid (TAG) band intensity was quantified using Fiji ImageJ software, and lipid concentrations were calculated from the standard curve generated with standard mixture. In the case of dissected larval guts (20-30 mid-guts per biological replicate, n = 3) dried lipids for all samples were re-suspended in 120 µl of chloroform and final lipid concentrations were calculated by normalizing protein in sample, measured by standard Bradford assay (BioRad #5000006).

### Adult starvation assay

Freshly emerging adult flies were collected over a period of 1 to 3 days and transferred to fresh standard food bottles. Flies were aged for 7 to 10 days, then male and female flies (100 each) were separately moved to food-free vials containing a cotton plug at the bottom soaked with 2 ml of tap water. Flies were kept at 25°C and vials were examined at least 3X daily to record number of dead flies. Surviving flies were moved to fresh vials with water-soaked plug every 24 hours for optimal hydration and to avoid fungal growth.

### Adult lifespan assay

Flies hatching between 0 to 3 hours were pooled, and males and females (30 each) were separately to fresh food. Vials were maintained at 25°C, and dead flies were scored daily while moving surviving flies to fresh food.

### Adult survivorship assay

Double FB-KDs of FASN1 and candidate genes were generated by crossing 5 to 7 *Dcg-Gal4>UAS-FASN1-RNAi* males to 20-25 females from candidate *UAS-RNAi* line in standard food vials cultured at 25°C. Mated females were allowed to lay eggs for 2 days, then parents were removed to fresh vials. L3 larvae carrying *Dcg-Gal4* driver + both *UAS-RNAi* transgenes were identified and moved to a new vial for developmental monitoring. Animals successfully completing metamorphosis and hatching into adult flies were scored. This count was compared to adult survivorship of *Dcg-Gal4* control, *Dcg-Gal4>UAS-FASN1-RNAi*, *Dcg-Gal4>cand-UAS-RNAi* animals.

### Fertility assays

All parents (*Dcg-Gal4* and *UAS-FASN1-RNAi* males and females) used were 7 to 10 days old. 10 virgin females were crossed with 10 males in fresh food vials streaked with yeast paste and kept overnight at 25°C. The following morning, flies were flipped to new food at 1 h intervals (3X) to empty females of older eggs/embryos. Flies were then allowed to lay on fresh plates supplemented with a drop of yeast paste for 4-6 h, following which all parents were removed from bottles and number of eggs laid was counted. The hatching of eggs on the food surface and development of progeny was closely monitored and recorded.

### Sample preparation for proteomic profiling

Fat bodies were extracted from late feeding L3 larvae (60-80 late-feeding L3 larvae per biological replicate, n = 3) in chilled Lysis Buffer + protease inhibitor cocktail. Tissues were homogenized by sonication and supernatant (lysate) was collected after centrifugation for 30 sec at 1000xg at 4°C. Centrifugation was repeated at 16,000xg for 10 min at 4°C, and supernatant transferred to fresh Eppendorf tube. Sample for loading was prepared by mixing protein sample with 2X LDS Sample Buffer and Lysis Buffer, and boiling at 95°C for 5 min. Sample was run on a 10% pre-cast polyacrylamide gel (BioRad) at 100 V for ∼25 min until the dye front was past the ladder. Gel was rinsed in MQ-water, then stained with Coomassie BB R250 in 10% acetic acid, 50% methanol, 40% water for 20 min at room temperature with rocking. Subsequently, gel was de-stained for 1 hour in 10% acetic acid, 50% methanol, 40% water with 4 solvent changes. When gel background appeared clear, de-stain was exchanged for water. Protein gel bands were cut on clean glass surface of gel imager with new razor, diced into 1 mm cubes, and moved to sterile Eppendorf tubes on ice for delivery to Proteomics Core Facility at UTSW for protein identification by mass spectrometry.

### FB tissue immunofluorescence

FBs dissected from late feeding L3 larvae were fixed in 4% paraformaldehyde at room temperature for 30 min on a nutator, then washed four times with 1 X PBST (PBS in 0.1% Tween), 10 min each. Tissues were blocked in 1 X PBST +2% normal goat serum for 2 h at room temperature, then incubated with anti-GFP recombinant rabbit monoclonal primary antibody (1:250 dilution, Thermo Fisher #G10362) added to the blocking buffer, overnight at 4°C. Next, FBs were washed 4X with 1 X PBST 15 min each, followed by incubation with goat anti-rabbit Alexa546 secondary antibody (1:1000 dilution, Invitrogen #A11035) in blocking buffer at room temperature in the dark for 2 h on a nutator. FBs were rinsed again four times in 1 X PBST 15 min each and subsequently mounted in SlowFade Gold antifade reagent with DAPI. Fluorescent structures were observed and imaged by confocal microscopy using standard RFP channel filter.

## Supplemental Figures

**SFigure 1:** A) Schematic of the steps of *de novo* lipogenesis (DNL), illustrating the key enzymes and metabolic products involved. B) Buoyancy-based test of fat content on extracted FBs from control and *FASN1^FB-RNAi^* larvae. C) Confocal images of FB periphery and mid-plane sections from *Dcg-Gal4* control alongside FASN loss-of-functions paralogs (*FASN1^FB-RNAi^*, *FASN2* global mutant, *FASN3^FB-RNAi^*), stained with LD stain monodansylpentane (MDH). Scale bar 50 µm. D) Confocal images of FB mid-plane sections from *Dcg-Gal4*, *Cg-Gal4*, and *r4-Gal4* driver control alongside their respective *FASN1* loss-of-function counterparts, stained with LD stain MDH. Scale bar 50 µm. E) Relative triglycerides (TG) in isolated larval guts from *Dcg-Gal4* control and *FASN1^FB-RNAi^*L3 larvae. Data are shown as mean ± SD from 3 biological replicates per genotype (20-30 guts per replicate), normalized to protein. P values were determined using an unpaired, two-tailed Welch’s t-test.

**SFigure 2:** A) Fertility assay performed to test viability of eggs produced by control (*Dcg-Gal4*) or *FASN1^FB-RNAi^* females mated with either *Dcg-Gal4* or *FASN1^FB-RNAi^* males. B) Confocal images of ovaries from 7–10-day old *Dcg-Gal4* control and *FASN1^FB-RNAi^* adult females stained with CellMask Actin stain. Entire *FASN1^FB-RNAi^*ovary (right) is visible at 10X magnification. Parts of the control ovary are labelled (left). Scale bar 100 µm. C) Confocal images of phalloidin-stained ovaries from control and *FASN1^FB-RNAi^* females. White arrows point to mature oocyte in control ovary and arrested oocyte in *FASN1^FB-RNAi^*. Scale bar (10X) 100 µm, Scale bar (20X) 50 µm. D) Confocal images of ovaries from *Dcg-Gal4* control and *FASN1^FB-RNAi^* females stained with CellMask Actin stain. Yellow arrows point to follicle cells of stage 9/10 egg chambers. White arrows point to oocyte; oocyte maturation fails in *FASN1^FB-RNAi^* ovaries. Scale bar 50 µm. E) Confocal images of LDs in control and *FASN1^FB-RNAi^* ovaries stained with BODIPY. Orange arrow in control ovary indicates LDs accumulated in nurse cells and purple arrow shows ring canal where movement of LDs from nurse cells to the developing oocyte is occurring. There also appear to be plentiful LDs in the mature oocyte*. LDs are sparse in *FASN1^FB-RNAi^* egg chambers. Scale bar (10X) 100 µm. Scale bar (20X) 50 µm F) Confocal images of ovaries from control and *FASN1^FB-RNAi^* females stained with nuclear stain DAPI. Blue arrow in control ovary point to distinct nuclei in mature oocytes, absent in *FASN1^FB-RNAi^* ovary. Scale bar (10X) 100 µm, Scale bar (20X) 50 µm.

**SFigure 3:** A) Coomassie dye extracted from guts of late-feeding L3 larvae, quantified as a measure of food intake. Data are shown as mean ± SD from 3 biological replicates per genotype (30 larvae per replicate). P values were determined using Welch’s one-way ANOVA. B) qPCR expression profiles of genes related to glycogenesis and glycogenolysis in FBs from *Dcg-Gal4* control and *FASN1^FB-RNAi^*. Data are presented as mean ± SD from 3 biological replicates per genotype (FBs pooled from 20-30 L3 larvae per replicate). P values were determined using an unpaired, two-tailed Student’s t-test. C) Glycogen levels measured in *Dcg-Gal4* control and *FASN1^FB-RNAi^* animals during metamorphosis: Feeding Larvae FL (∼100-108 h AEL), Pharate Adults PA (∼205-215 AEL), AEL = After Egg Laying. Data are shown as mean ± SD from 3 biological replicates per genotype (10 animals per replicate), normalized to weight. P values were determined using Welch’s one-way ANOVA. D) TG levels measured in *Dcg-Gal4* control and *FASN1^FB-RNAi^* animals during metamorphosis: Feeding Larvae FL (∼100-108 h AEL), Pharate Adults PA (∼205-215 AEL), AEL = After Egg Laying. TLC plate representing TG bands corresponding to the quantified samples is shown below the graph (sample volumes were normalized to larval weight). Data are shown as mean ± SD from 3 biological replicates per genotype (10 animals per replicate), normalized to weight. P values were determined using Welch’s one-way ANOVA. E) Glycogen measured in *Dcg-Gal4* control and *FASN1^FB-RNAi^* male and female flies. Data are shown as mean ± SD from 3 biological replicates per genotype, per gender (10 flies per replicate). P values were determined using Welch’s one-way ANOVA.

**SFigure 4:** A) Confocal section images of FB periphery and mid-plane stained with LD stain MDH. FBs from the ubiquitous but mosaic driver *da-Gal4* (control) and *da-Gal4>UAS-FASN1^RNAi^* L3 larvae also express PM marker mcd8-GFP; green indicates where *da-gal4* is expressed, and where *FASN1-*KD also occurs (right only). Scale bar 50 µm. B) PAS glycogen stain in FBs from driver *da-Gal4* and *da-Gal4>UAS-FASN1^RNAi^*. White arrows indicate where *da-gal4* is expressed and *FASN1-*KD occurs (stronger PAS stain). Blue arrows point to regions where *FASN1* KD is inactive. Scale bar 50µm. C) Confocal images of FBs from RNAi lines targeting DNL enzymes, stained with LD stain MDH. Scale Bar 25 µm. D) PAS glycogen staining of FBs from RNAi lines targeting DNL enzymes. Scale Bar 50 µm. E) Relative change in SREBP protein levels in *FASN1^FB-RNAi^*FBs detected by global proteomics. Data are shown as mean ± SD from 3 biological replicates per genotype (60-80 FBs per replicate). P values were determined using an unpaired, two-tailed Student’s t-test.

**SFigure 5:** A) qPCR analyses in FBs from *FASN1^FB-RNAi^* and *SREBP^FB-RNAi^* single KD and *FASN1^FB-RNAi^* + *SREBP^FB-RNAi^* KD double L3 larvae to validate depletion of *FASN1*and *SREBP*. Data are shown as mean ± SD from 3 biological replicates per g notype (FBs pooled from 20-30 larvae per replicate). P values were determined using an unpaired, two-tailed Student’s t-test. B) Confocal images of mid-plane sections of FBs from *FASN1^FB-RNAi^* and *SREBP^FB-RNAi^* single KD and *FASN1^FB-RNAi^* + *SREBP^FB-RNAi^*double KD L3 larvae stained with LD stain MDH. Scale Bar 50 µm. C) Total triglycerides (TG) levels measured by thin layer chromatography (TLC) in whole *Dcg-Gal4*, *FASN1^FB^*^-RNAi^, *SREBP^FB-RNAi^*, and *FASN1^FB-RNAi^* + *SREBP^FB-RNAi^* L3 larvae. TLC plate representing TG bands corresponding to the quantified samples is shown below the graph (sample volumes loaded onto the plate were normalized to larval weight such that band intensities reflect TG abundance). Data are shown as mean ± SD from 3 biological replicates per genotype (15 larvae per replicate). P values were determined using Welch’s one-way ANOVA. D) Relative expression of *tobi* mRNA in FB from *Dcg-Gal4*, *FASN1^FB^*^-RNAi^, *SREBP^FB-RNAi^*, and *FASN1^FB-RNAi^* + *SREBP^FB-RNAi^* larvae. Data are shown as mean ± SD from 3 biological replicates per genotype (FBs pooled from 20-30 larvae per replicate). P values were determined using an unpaired, two-tailed Student’s t-test.

## Supplemental Tables

**Table S1:** List of Drosophila stock lines used in this study.

**Table S2**: Mass spectrometry proteomics dataset for isolated larval fat bodies. The FASN1 larval fat body specific RNAi knockdown is compared to Dcg-Gal4 control fat bodies. The three FASN1 KD repeats are listed on sheet two as FASN 1-3. Dcg-Gal4 control fat bodies are Dcg 1-3.

