## Supplementary material for "SREBP regulates a cell autonomous triglyceride:glycogen metabolic switch in *Drosophila*": Supp Figures

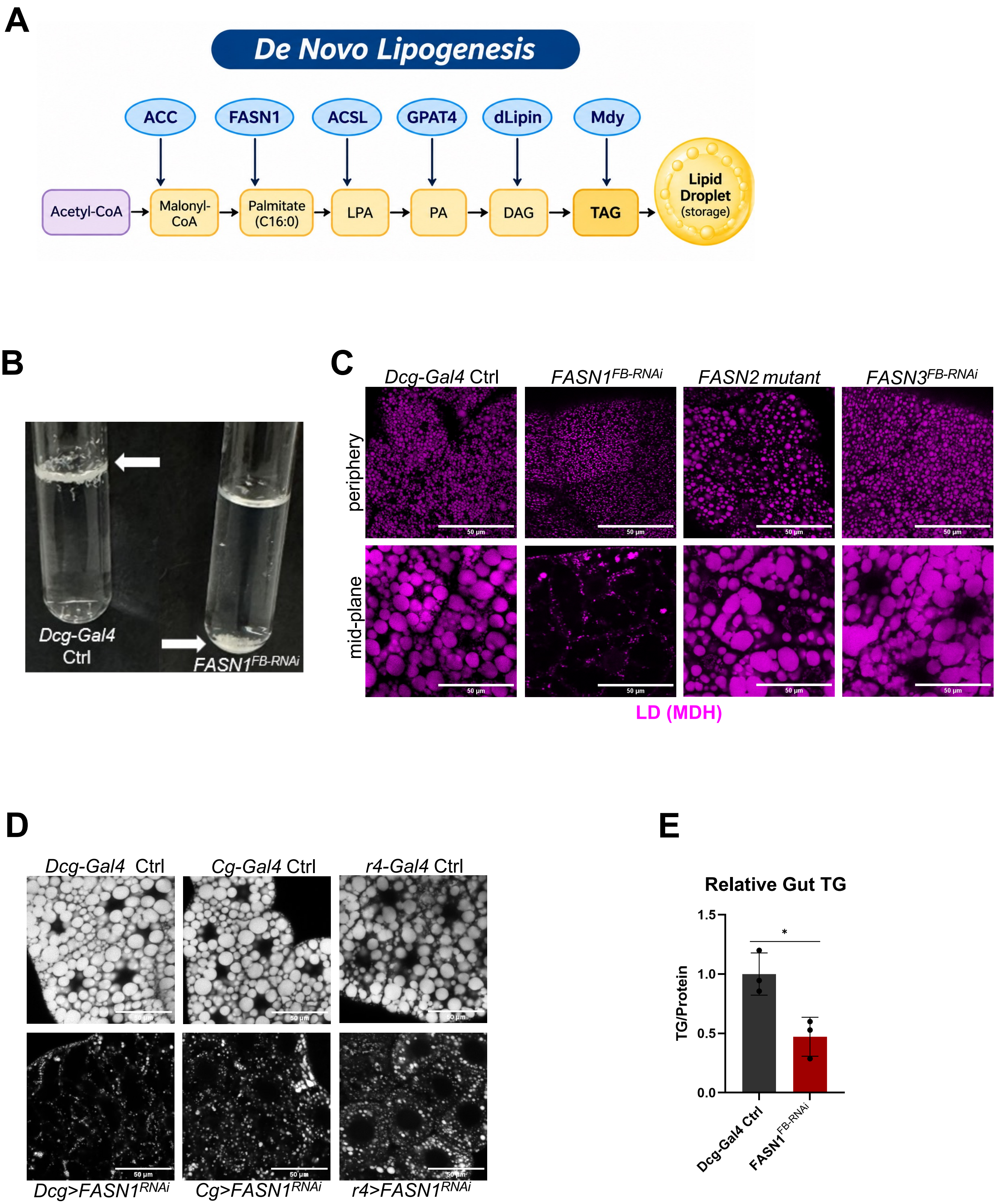

A
De Novo Lipogenesis
ACC
FASN1
ACSL
GPAT4
dLipin
Mdy
Acetyl-CoA
Malonyl-CoA
Palmitate (C16:0)
LPA
PA
DAG
TAG
Lipid Droplet (storage)
B
Dcg-Gal4 Ctrl
FASN1FB-RNAi
C
Dcg-Gal4 Ctrl
FASN1FB-RNAi
FASN2 mutant
FASN3FB-RNAi
periphery
mid-plane
50 μm
LD (MDH)
D
Dcg-Gal4 Ctrl
Cg-Gal4 Ctrl
r4-Gal4 Ctrl
Dcg>FASN1RNAi
Cg>FASN1RNAi
r4>FASN1RNAi
E
Relative Gut TG
TG/Protein
1.5
1.0
0.5
0.0
*
Dcg-Gal4 Ctrl
FASN1FB-RNAi


S Figure 1

A

**Fertility Assay**

| Males | Females | #Eggs | #Adults | % Survival |
|---|---|---|---|---|
| *Dcg-Gal4* | *Dcg-Gal4* | 415 | 382 | 93 |
| *FASN1*$^{FB\text{-}RNAi}$ | *Dcg-Gal4* | 416 | 288 | 69 |
| *Dcg-Gal4* | *FASN1*$^{FB\text{-}RNAi}$ | 3 | 0 | |
| *FASN1*$^{FB\text{-}RNAi}$ | *FASN1*$^{FB\text{-}RNAi}$ | 4 | 3 | |

B

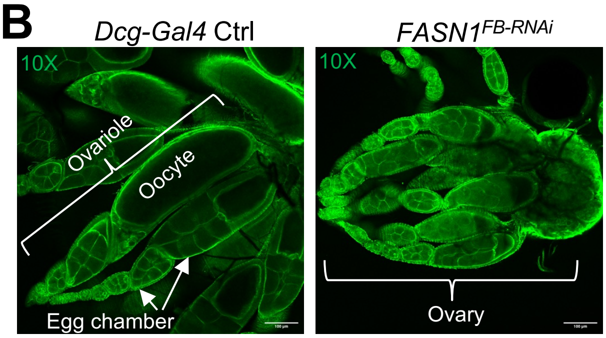


CellMask Actin stain

C

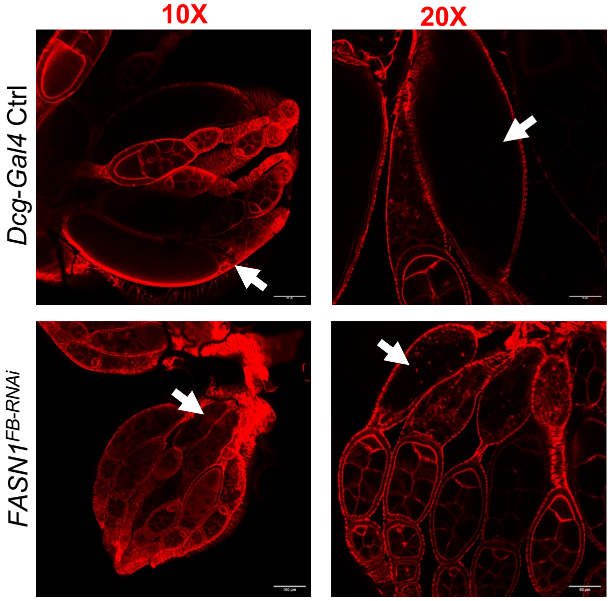


Phalloidin

D E F

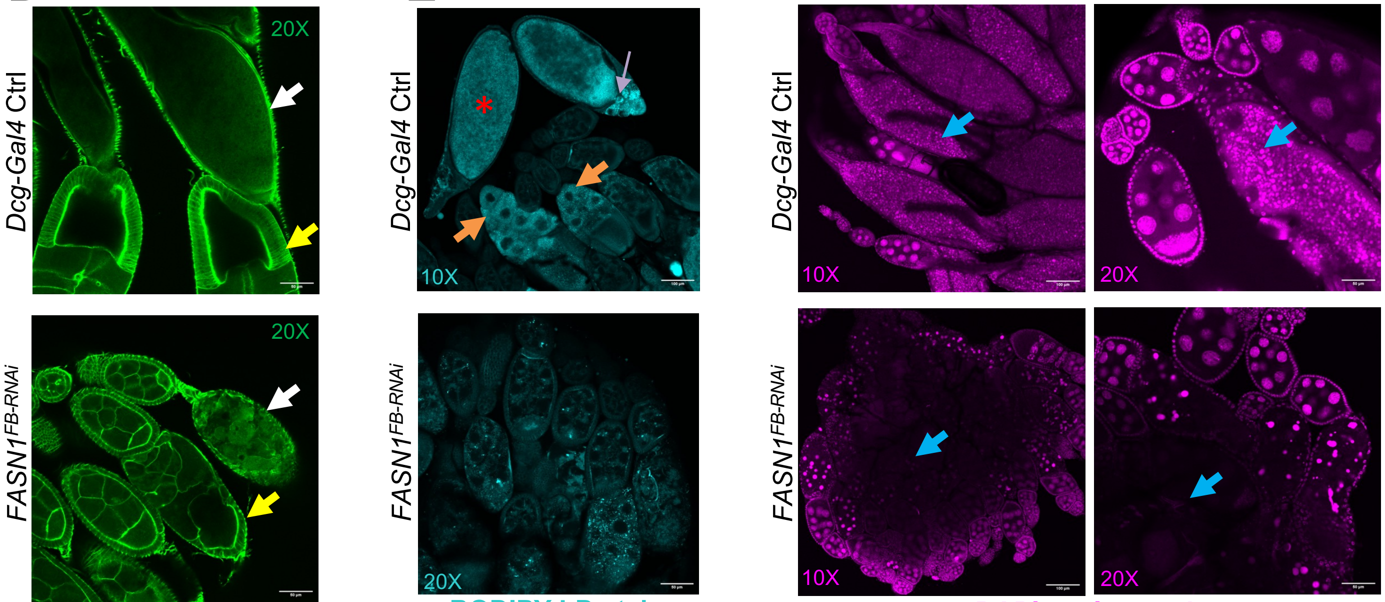


CellMask Actin stain

BODIPY LD stain

DAPI nuclear stain

**Supp Fig. 2**

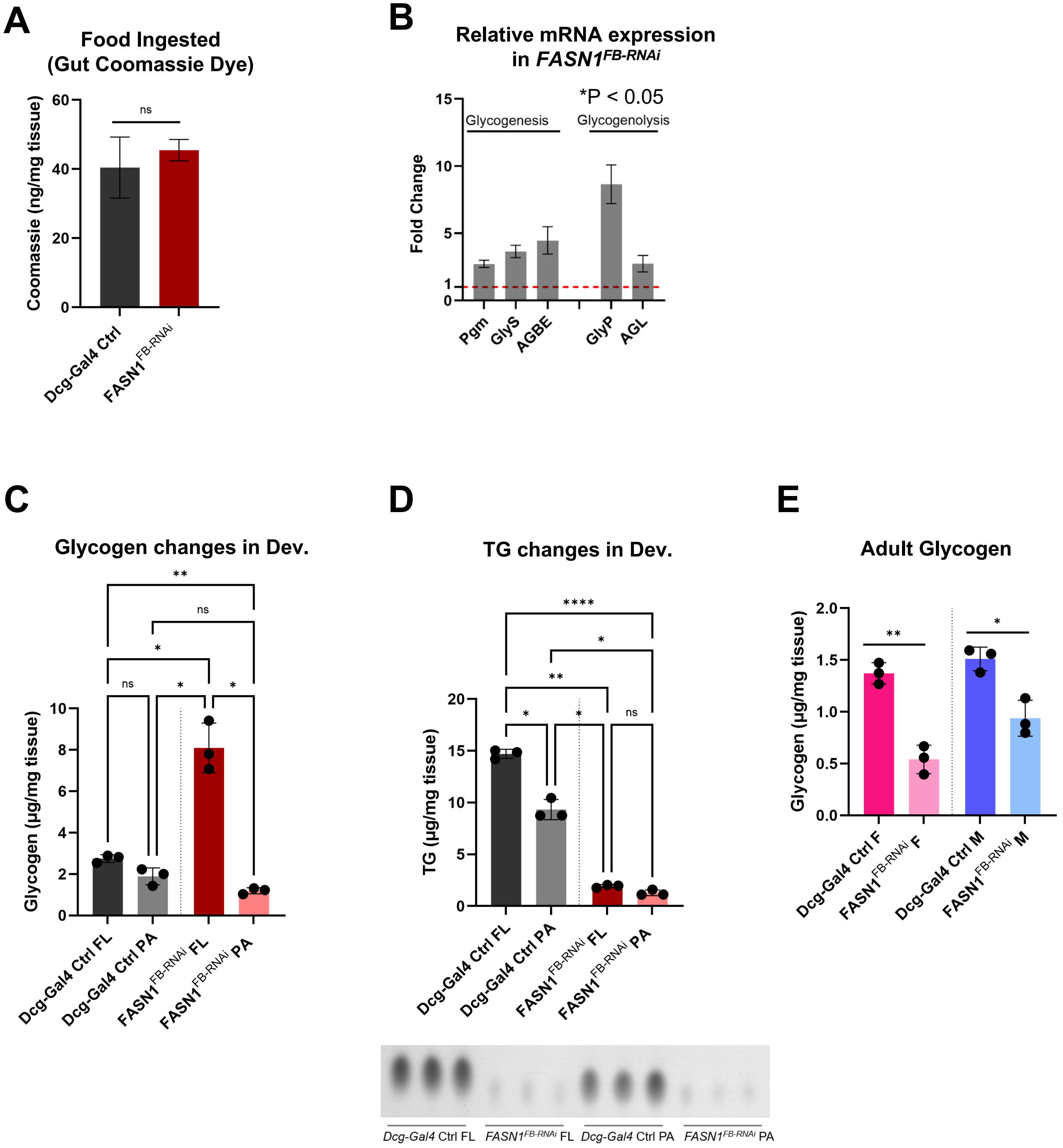

A
Food Ingested (Gut Coomassie Dye)
Coomassie (ng/mg tissue)
60
40
20
0
ns
Dcg-Gal4 Ctrl
FASN1 FB-RNAi
B
Relative mRNA expression in FASN1 FB-RNAi
*P < 0.05
Fold Change
15
10
5
1
0
Glycogenesis
Glycogenolysis
Pgm
GlyS
AGBE
GlyP
AGL
C
Glycogen changes in Dev.
Glycogen (µg/mg tissue)
10
8
6
4
2
0
**
ns
*
ns
*
*
Dcg-Gal4 Ctrl FL
Dcg-Gal4 Ctrl PA
FASN1 FB-RNAi FL
FASN1 FB-RNAi PA
D
TG changes in Dev.
TG (µg/mg tissue)
20
15
10
5
0
****
*
**
*
*
ns
Dcg-Gal4 Ctrl FL
Dcg-Gal4 Ctrl PA
FASN1 FB-RNAi FL
FASN1 FB-RNAi PA
Dcg-Gal4 Ctrl FL
FASN1 FB-RNAi FL
Dcg-Gal4 Ctrl PA
FASN1 FB-RNAi PA
E
Adult Glycogen
Glycogen (µg/mg tissue)
2.0
1.5
1.0
0.5
0.0
**
*
Dcg-Gal4 Ctrl F
FASN1 FB-RNAi F
Dcg-Gal4 Ctrl M
FASN1 FB-RNAi M


**S Figure 3**

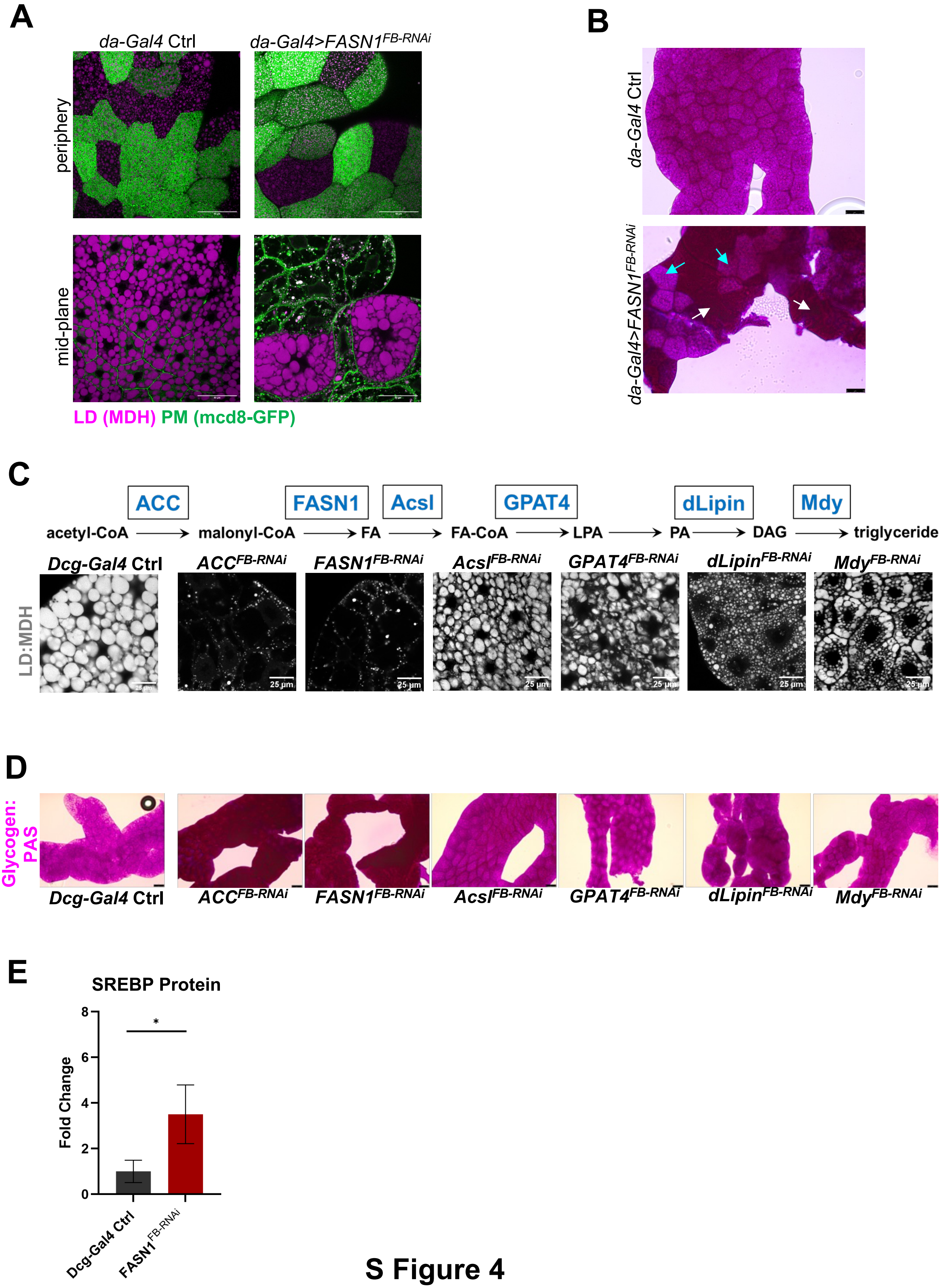
A
da-Gal4 Ctrl
da-Gal4>FASN1FB-RNAi
periphery
mid-plane
LD (MDH) PM (mcd8-GFP)
B
da-Gal4 Ctrl
da-Gal4>FASN1FB-RNAi
C
ACC
FASN1
Acsl
GPAT4
dLipin
Mdy
acetyl-CoA
malonyl-CoA
FA
FA-CoA
LPA
PA
DAG
triglyceride
Dcg-Gal4 Ctrl
ACCFB-RNAi
FASN1FB-RNAi
AcslFB-RNAi
GPAT4FB-RNAi
dLipinFB-RNAi
MdyFB-RNAi
LD:MDH
25 µm
D
Glycogen: PAS
Dcg-Gal4 Ctrl
ACCFB-RNAi
FASN1FB-RNAi
AcslFB-RNAi
GPAT4FB-RNAi
dLipinFB-RNAi
MdyFB-RNAi
E
SREBP Protein
Fold Change
0
2
4
6
8
*
Dcg-Gal4 Ctrl
FASN1FB-RNAi

S Figure 4

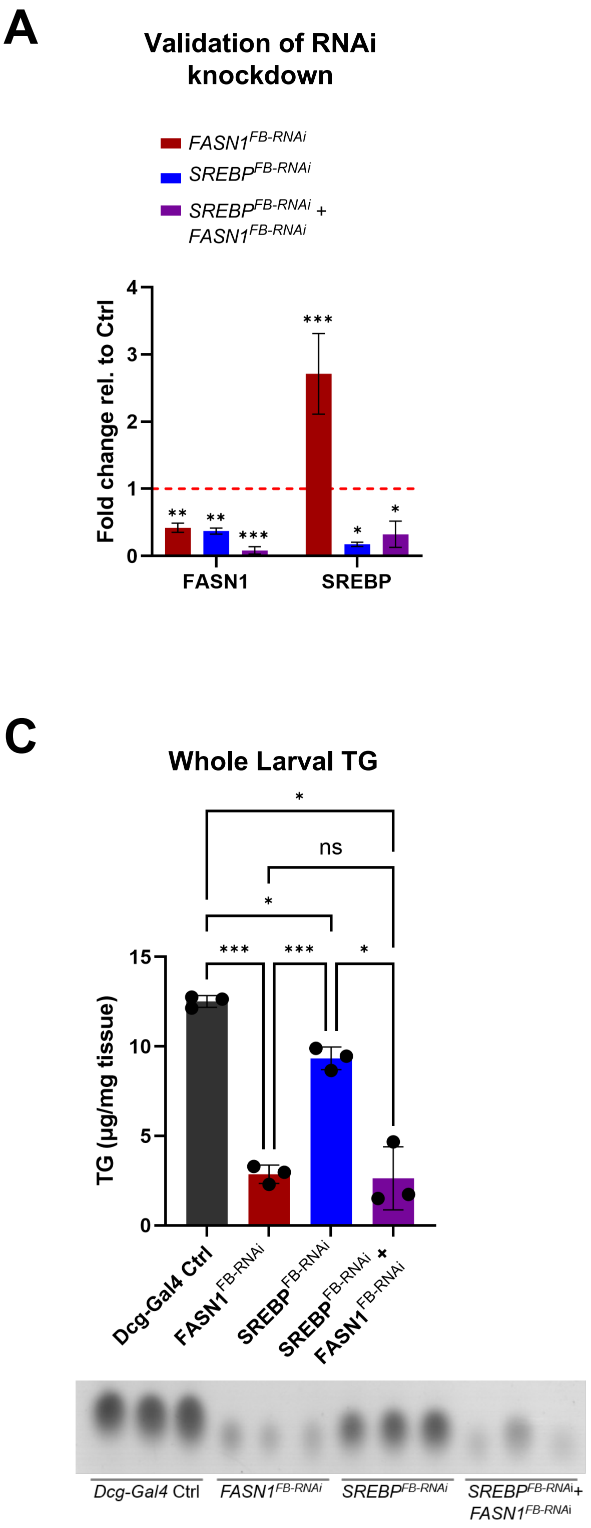

A
Validation of RNAi knockdown
FASN1FB-RNAi
SREBPFB-RNAi
SREBPFB-RNAi + FASN1FB-RNAi
Fold change rel. to Ctrl
FASN1
SREBP
B
Dcg-Gal4 Ctrl
FASN1FB-RNAi
SREBPFB-RNAi
SREBPFB-RNAi +FASN1FB-RNAi
50 µm
C
Whole Larval TG
TG (µg/mg tissue)
Dcg-Gal4 Ctrl
FASN1FB-RNAi
SREBPFB-RNAi
SREBPFB-RNAi + FASN1FB-RNAi
D
tobi mRNA
Fold Change


### S Figure 5
